# A minor muscle stem cell population functions as metabolic Trojan horse fueling systemic aging by succinate induced epigenetic reprogramming

**DOI:** 10.64898/2025.12.26.696560

**Authors:** Xiaofen Wu, Meng Chen, Ruqi Fan, Hongye Wang, Ruimiao Ma, Xiaocui Tang, Hui Wang, Wei Wei, Jiemin Wong, Guoliang Xu, Ping Hu

## Abstract

Inflammaging, the sustained chronic inflammation, is a hallmark of aging, yet its sustained activation mechanism remains elusive. Here, we identified muscle stem cells (MuSCs) as a driver of systemic inflammaging, evidenced by the multi-organ inflammation and aging phenotypes in MuSC specific Tet2 knockout mice. Tet2-Hdac11-Acod1-SDH axis maintained normal succinate level in MuSCs. Tet2 knockout disrupted this enzymatic cascade and led to succinate accumulation, fueling H4K31succ elevation to directly activate inflammatory gene transcription in MuSCs. The excess succinate was delivered to muscle fibers by sporadic fusion of Tet2 knockout MuSCs during muscle homeostasis, activated pro-inflammatory program, and transformed muscle to a persistent pro-inflammatory factor secretory organ, sustaining systemic inflammaging. Moreover, Tet2 was downregulated in aged MuSCs suggesting that this coupled metabolic-epigenetic mechanism was active in physiological aging. These findings reveal that a small subset of dysregulated MuSCs activate sustained whole-body inflammaging and multi-organ aging, providing new targets for rejuvenation strategy development.

## INTRODUCTION

Ten-Eleven translocation 2 (Tet2) is a key epigenetic regulator belonging to the Tet DNA dioxygenase family that catalyzes the iterative oxidation of 5-methylcytosine (5mC) to 5-hydroxymethylcytosine (5hmC), 5-formylcytosine (5fC), and 5-carboxylcytosine (5caC). This series of reactions result in active DNA demethylation, as 5caC is excised by thymine DNA glycosylase (TDG) and replaced with unmodified cytosine via base excision repair.^1^ Tet2 plays critical roles in many cellular processes, such as the maintenance of naïve pluripotency, immune cell development, and tumorigenesis.^2,3^ We have previously shown that *Tet2* knockout (KO) leads to impaired muscle regeneration after acute injury. Tet2 facilitates MyoD binding by maintaining hypomethylation around MyoD recognition sites to promote myogenesis.^4^ DNA methylation is a well-established aging clock.^5^ As an enzyme playing central role in regulating DNA methylation, Tet2 has been shown to regulate organ function changes during aging. The up-regulation of *Tet1* and *Tet2* has been shown to counteract age related vision loss in mice by restoring youthful DNA methylation patterns.^6^ The expansion of hematopoietic cell clones containing somatic *Tet2* mutations during aging increases the risk of atherosclerosis.^7,8^ Tet2 has been reported to regulate bone aging and bone marrow stromal cell metabolism.^9^ These findings underscore the multifaceted role of Tet2 in aging, warranting further investigations into the mechanism of the functions of it across different tissues.

Inflammaging, a sustained chronic, sterile, and low-grade inflammation developed during aging, is a key hallmark of aging which increases the risk for many age-related chronic diseases such as chronic obstructive pulmonary disease (COPD), lung fibrosis, and liver fibrosis.^10-12^ The mechanism driving inflammaging involves a complex interplay of multiple elements. Senescence associated secretory phenotype (SASP), comprising a mixture of pro-inflammatory factors including cytokines, chemokines, and other factors secreted by senescent cells, has been considered to be one of the drivers of inflammaging.^13^ The accumulation of senescent cells and SASP are associated with increased inflammation in both mice and human tissues.^14^ Damage associated molecular patterns (DAMPs), that stimulate TLR/cGAS-STING and NFκB signaling, are also attributed to inflammaging during aging.^15,16^ Despite of these insights, the mechanism governing sustained inflammaging during aging remains to be fully elucidated.

Metabolites generated by central carbon metabolism provide not only energy to support cellular activity but also substrates or cofactors for epigenetic enzymes, thereby linking metabolic state to gene expression.^17,18^ Tricarboxylic acid (TCA) cycle is one of the key interface of the metabolism-epigenetic crosstalk. For instance, Acetyl-CoA is an essential substrate for histone lysine acetyltransferases (KATs).^19^ Similarly, succinate, a TCA cycle intermediate metabolite, regulates cellular activities by multiple means. It is a substrate for SDH, also known as mitochondria Complex II.^20^ SDH has been shown activating T cell proliferation and repressing inflammation.^21^ Loss of function mutation of SDH is associated with tumorigenesis.^22^ Succinate accumulation promotes tumorigenesis and delays pluripotent stem cell differentiation by altering epigenetic landscape.^18,23,24^ Due to the structure similarity, succinate competitively inhibits the activities of α-ketoglutarate (α-KG) dependent JMJD family histone demethylases and TET DNA demethylases.^23^ Succinate can also serve as a substrate for protein lysine succinylation and regulate protein functions.^25^ Succinate accumulation has been shown to promote inflammation.^26,27^ During aging, defective mitochondria and declined SDH activity are observed in muscle cells.^28-30^ Supplementation of α-KG, which is considered to be an antagonist of succinate in some respect,^23,31^ improves proliferation of MuSCs and muscle regeneration in aged mice.^32^ However, the mechanism governing the functions of succinate during aging still needs further investigation.

Skeletal muscle is the most abundant tissue in human body, constituting approximately 40% of the total body mass.^33^ The maintenance and regeneration of muscle throughout postnatal life are primarily driven by MuSCs. Under homeostatic conditions, most MuSCs remain in a quiescent state in their niche. In response to muscle growth or injury, quiescent MuSCs are rapidly activated and proliferate, generating sufficient cells to fuse to existed myofibers or form new myofibers to facilitate tissue reparation or growth.^34-36^ During cell fusion, MuSCs contribute not only nuclei but also mitochondria and cellular metabolites to myofibers.^37^ The functional significance of these materials donated by MuSCs was not clear yet.

Muscle mass and function progressively decline with age, which eventually develops to sarcopenia. Decreased respiratory and metabolic functions are usually accompanied with sarcopenia. Multiple factors including protein catabolism, reduced protein anabolism, alteration of hormone levels, and malnutrition are considered to be the primary driver for muscle aging and sarcopenia.^38,39^ MuSCs have undergone a spectrum of age-related changes such as loss of quiescence, increased senescence, greater heterogeneity, and declined differentiation ability.^40^ The decline of MuSC function is considered mainly contributing to the diminished muscle regenerative potential upon injury in aged population.^30,32,41-43^ Under the non-injury homeostatic condition, the link between MuSCs and organ aging has not been fully understood.

Here, we demonstrate that MuSC specific ablation of *Tet2* led to systemic inflammation, multi-organ fibrosis, and premature aging. Further analysis revealed Tet2-Hdac11-Acod1-SDH axis of enzymatic reactions controlling cellular succinate level. In *Tet2* KO MuSCs, the aberrant succinate accumulation increased H4K31 succinylation, altering the epigenetic landscape to activate inflammatory gene transcription. The excess succinate in Tet2 KO MuSCs was carried to myofibers through cell fusion during muscle homeostasis. In myofibers, succinate similarly elevated H4K31succ level and drove inflammatory gene expression, thereby transforming skeletal muscle into a potent source for inflammatory signaling that sustained systemic inflammation, accelerated multi-organ fibrosis and aging. Notably, *Tet2* expression was significantly down-regulated and succinate level was up-regulated in aged MuSCs, and the same succinate mediated inflammation amplification mechanism was active during natural aging. Importantly, citraconate treatment, which reduced the cellular succinate level, reversed the aging phenotypes, suggesting that targeting succinate directed metabolic reprogramming is a potential rejuvenation strategy.

## RESULTS

### *Tet2* loss in MuSCs results in premature multi-organ aging

We have previously demonstrated that *Tet2* KO in MuSCs impaired muscle regeneration after acute injury in young mice.^4^ To investigate the long-term consequences of *Tet2* ablation, MuSC specific *Tet2* KO was induced by Tamoxifen (TAM) injection in 4-week-old *Pax7* CreERT2: *Tet2* flox/flox mice (Figure 1A). Consistent with our previous report,^4^ *Tet2* KO mice were largely normal in body size, appearance, and muscle size 2 weeks after *Tet2* KO induction (6 weeks of age) (Figures S1A-E). However, 3 months after *Tet2* KO in MuSCs (16 weeks of age), premature aging phenotypes were observed as indicated by reduced body size and weight, severe hair loss, and a high frequency of hunched backs (Figures 1B-E). To determine whether these appearances were correlated with aging related functional and structure decline in multiple organs, skeletal muscle, lung, and liver were further analyzed. *Tet2* KO mice showed a decline in muscular motor functions, evidenced by reduced running distance and time on treadmills (Figure 1F). This was accompanied by decreased muscle weight and muscle fiber size (Figures 1G and H). In situ muscle force test indicated that both the twitch and tetanic force were decreased in *Tet2* KO mice (Figure 1I). And increased muscle fatigue was also observed in *Tet2* KO mice (Figure 1J). Together, these phenotypes, which mimic sarcopenia, suggest that *Tet2* KO in MuSCs drives premature aging of skeletal muscle.

**Figure 1.**
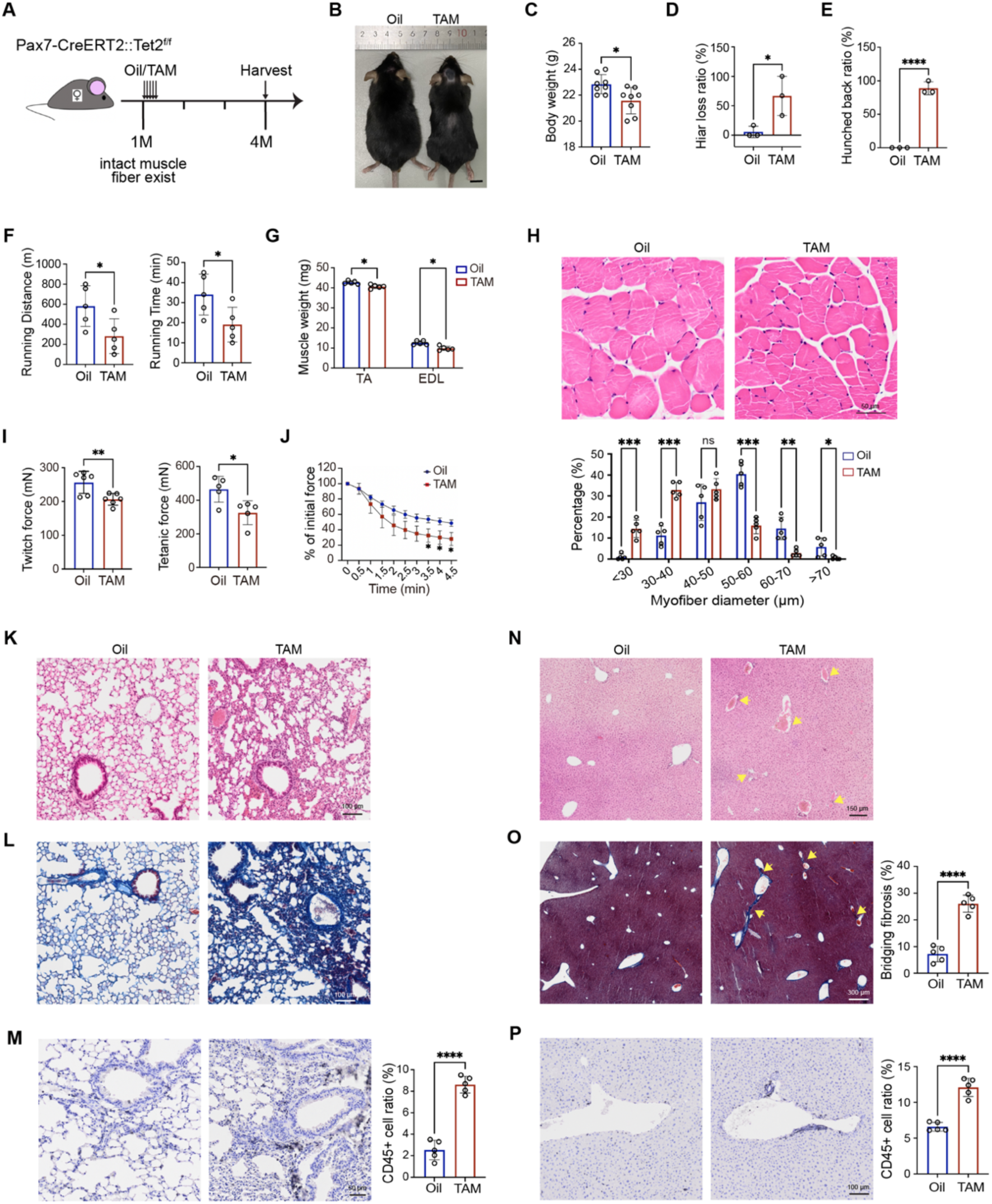
MuSC specific knockout of *Tet2* leads to multi-organ aging. **(A)** Scheme of induction of MuSC specific knockout of *Tet2*. Either oil (Oil) or tamoxifen (TAM) was injected to *Pax7* CreERT2:*Tet2* f/f mice at the age of 1-month-old (1M). Samples were harvested 3 months after TAM injection (4-month-old, 4M) for functional assays. **(B)** Representative images of 4-month-old MuSC specific *Tet2* KO mice. Scale bar: 1cm. **(C)** Statistical analysis of body weights of 4-month-ol Oil (f/f) or TAM injected (Tet2 KO) mice (n = 8 in each group). **(D)** Statistical analysis of hair loss ratio in 4-month-old Oil (f/f) or TAM injected (*Tet2* KO) mice (n = 18 in each group). The ratio was calculated based on data collected from 3 batches of mice, 6 mice in each batch. **(E)** Statistical analysis of hunched back ratio of in 4-month-old Oil (f/f) or TAM injected (*Tet2* KO) mice (n = 18 in each group). The ratio was calculated based on data collected from 3 batches of mice, 6 mice in each batch. **(F)** Mean running time and running distance of 4-month-old Oil (f/f) or TAM injected (*Tet2* KO) mice when trained on treadmill (n = 5 in each group). **(G)** Extensor digitorum longus (EDL) and tibialis anterior (TA) muscle weights in 4-month-old Oil (f/f) or TAM injected (*Tet2* KO) mice (n = 5 in each group). **(H)** Representative images of H&E staining of TA muscles from 4-month-old Oil (f/f) or TAM injected (*Tet2* KO) mice (up), and percentage quantification of myofiber size (below). n=5 in each group. Scale bars, 50μm. **(I)** *In situ* Twitch and tetanic force of TA muscles from 4-month-old Oil (f/f) or TAM injected (*Tet2* KO) mice (n = 6 in each group). **(J)** *In situ* Fatigue characteristics of TA muscles from 4-month-old Oil (f/f) or TAM injected (*Tet2* KO) mice. Values at each time point indicated the percentage of the initial force at the beginning of the fatigue trial (time 0). **(K)-(L)** Representative images of H&E (K) and Masson (L) staining of lung from 4-month-old Oil (f/f) or TAM injected (*Tet2* KO) mice. Scale bars, 100μm. **(M)** Representative images of CD45 immunohistochemistry staining in lung from 4-month-old Oil (f/f) or TAM injected (*Tet2* KO) mice (left), and the quantification of the percentage of CD45-positive cells (right). n=5 in each group. Scale bars, 50μm. **(N)** Representative images of H&E staining of liver from 4-month-old Oil (f/f) or TAM injected (*Tet2* KO) mice. For H&E staining, area with potential immune cells infiltration was indicated by yellow arrows. Scale bars, 150μm. **(O)** Representative images of Masson staining of liver from 4-month-ol Oil (f/f) or TAM injected (*Tet2* KO) mice were shown at the left. Portal triads to Central vein (P-C) bridging fibrosis, which is a marker for severe liver fibrosis, was indicated by yellow arrows. Quantification of the percentage of bridging fibrosis was shown at the right. Scale bars, 300μm. **(P)** Representative images of CD45 immunohistochemistry staining in liver from 4-month-old Oil (f/f) or TAM injected (*Tet2* KO) mice (left), and the quantification of the percentage of CD45-positive cells (right). n=5 in each group. Scale bars, 100μm. **(C)-(J), (M) and (O)-(P)** Data were shown in mean ± SD form. Data were analyzed by unpaired t-test. * indicated *p* < 0.05; ** indicated *p* < 0.01; *** indicated *p* < 0.001.

Lung tissues were harvested from MuSC specific *Tet2* KO and WT mice. Hematoxylin and Eosin (H&E) and Masson staining were performed to evaluate the fibrosis level. Surprisingly, fibrosis was significantly increased in lung after *Tet2* KO in MuSCs (Figures 1K and L). Increased immune cell infiltration into the lung was also observed as indicated by H&E staining (Figure 1K), which was further confirmed by CD45 immunohistological staining (Figure 1M), suggesting elevated inflammation level.

Similarly, in the liver, increased fibrosis was observed as shown by H&E and Masson staining in *Tet2* KO mice (Figures 1N and O). A large amount of immune cell infiltration was also detected by CD45 immunohistological staining (Figure 1P), suggesting a concomitant rise in hepatic inflammation.

These unexpected results suggest that *Tet2* deficiency in MuSCs initiates a cascade of premature aging and the aging effect extended to multiple organs other than muscle by *Tet2* KO in MuSCs.

### *Tet2* KO leads to succinate accumulation in MuSCs

To elucidate the mechanism by which MuSC specific *Tet2* KO drives multi-organ defects, we first explored the cell intrinsic functions of Tet2 in MuSCs. Given the observed muscle fatigue in *Tet2* KO mice, a phenotype closely linked to mitochondria dysfunction,^44,45^ we went on to assess the mitochondria function in MuSCs. Mitotracker staining revealed that the length of mitochondria was significantly shorter in *Tet2* KO MuSCs (Figures 2A and B). The respiratory functions of mitochondria were further evaluated by seahorse assays. The basal oxygen consumption rate (OCR) of mitochondria from *Tet2* KO MuSCs was reduced and the ATP production decreased (Figures 2C and D), suggesting impaired mitochondria respiration resulted from *Tet2* KO in MuSCs.

**Figure 2.**
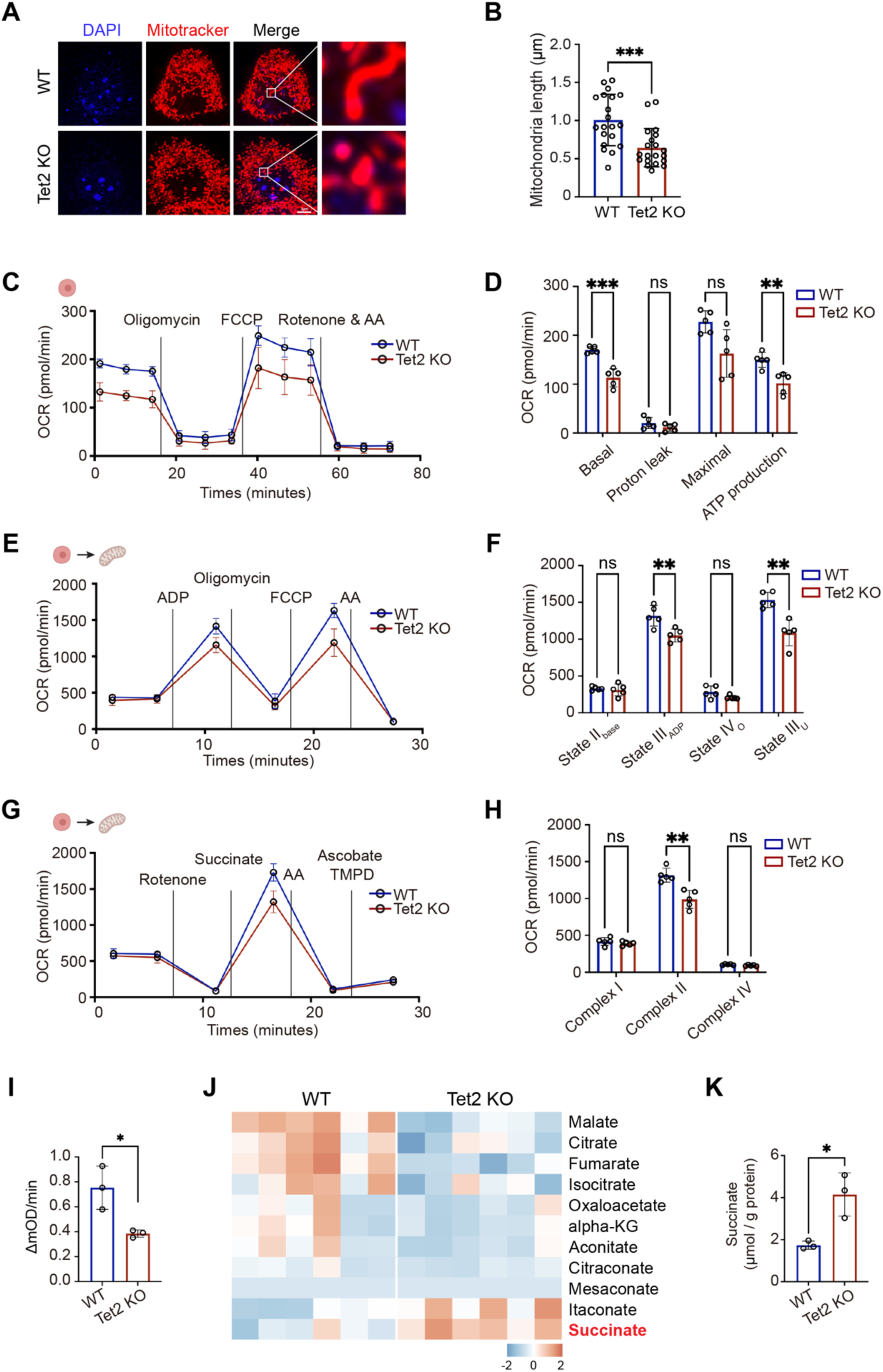
*Tet2* KO results in succinate accumulation in MuSCs. **(A)** Representative images of the mitochondria stained with MitoTracker (red) and DAPI (blue) in WT and *Tet2* KO MuSCs, respectively. Scale bars, 3μm. **(B)** Quantification of the mitochondria length from MitoTracker staining. **(C)** Seahorse assays showing normalized oxygen-consumption rates (OCRs) in WT and *Tet2* KO MuSCs, respectively. Data were normalized to cell numbers. n=5 in each group. **(D)** Quantification of normalized OCRs in basal respiration, proton leak, maximal respiration, and ATP production (n=5 in each group). **(E)** Seahorse coupling assays showing normalized OCRs in mitochondria isolated from WT and *Tet2* KO MuSCs, respectively. Data were normalized to protein level. n=5 in each group. **(F)** Quantification of normalized OCRs in state II_base_ (Basal respiration), state III_ADP_ (ADP-stimulated respiration), state IV_O_ (non-ADP-stimulated respiration, after oligomycin treatment), and state III_U_ (uncoupled respiration, after FCCP treatment) (n=5 in each group). **(G)** Seahorse respiratory complex assays showing normalized OCRs in mitochondria isolated from WT and *Tet2* KO MuSCs, respectively. Data were normalized to total protein level. n=5 in each group. **(H)** Quantification of respiration of each mitochondrial ETC complex (n = 5 in each group). **(I)** SDH enzyme activity measured by DCPIP assay in WT and *Tet2* KO MuSCs, respectively (n=3 in each group). **(J)** Heatmap of targeted LC-MS/MS metabolomics data in WT and *Tet2* KO MuSCs, respectively. The metabolites were color-coded to represent their relative levels in *Tet2* KO MuSCs compared to that in WT. **(K)** Absolute succinate content measured by coupled enzyme assay in WT and *Tet2* KO MuSCs, respectively (n=3 in each group). **(B)-(I) and (K)** Data were shown in mean ± SD form. Data were analyzed by unpaired t-test. * indicated *p* < 0.05; ** indicated *p* < 0.01; *** indicated *p* < 0.001.

To pinpoint the defective stage of electron transfer, substrate specific analysis was performed. The addition of ADP in seahorse assays failed to rescue respiration and the uncoupled State IIIu remained low (Figures 2E and F), suggesting dysfunctions in substrate oxidation and electron transfer chain (ETC). To further interrogate the function of ETC complexes, succinate was added to seahorse assays. The drop of OCR was observed, directly indicating Complex II (Succinate dehydrogenase, SDH) dysfunction (Figures 2G and H). Complex II (SDH), which is embedded in the TCA cycle catalyzing the conversion from succinate to fumarate, plays a critical role in succinate driven respiration. The compromised SDH activity was further confirmed by DCPIP (2, 6-dichlorophenol-indophenol) reduction assay (Figure 2I). Collectively, these results suggest that *Tet2* KO impairs complex II (SDH) function in MuSCs.

As SDH catalyzes the conversion of succinate to fumarate in TCA cycle, its dysfunction was expected to cause succinate accumulation. We therefore performed metabolome analysis using *Tet2* KO MuSCs. Indeed, succinate level increased in *Tet2* KO MuSCs (Figures 2J and S2A). To further validate the metabolome analysis results, whole cell extracts from *Tet2* KO MuSCs were subjected for succinate level assessment using coupled enzyme assay. The accumulation of succinate was confirmed (Figure 2K). Collectively, these results suggest that *Tet2* ablation in MuSCs compromises SDH activity, driving succinate accumulation, and metabolic dysregulation.

### Disruption of a Tet2-Hdac11-Acod1 cascade results in succinate accumulation in *Tet2* KO MuSCs

To define the mechanistic link between Tet2 and SDH activity, we first checked whether Tet2 regulated the expression of key metabolic enzymes catalyzing the synthesis and dehydrogenation of succinate. RNA-seq data showed that no changes in the expression of *Succinyl-CoA Synthetase* (*SCS*) and *SDH* isoforms (Figures S2B and C), suggesting that Tet2 does not directly regulate the expression of *SCS* and *SDH* isoforms.

Further analysis of the RNA-seq data revealed an up-regulation of *Acod1* in *Tet2* KO MuSCs (Figure S3A). Acod1 is an enzyme catalyzing the conversion of cis-aconitate to itaconate, which is a known inhibitor of SDH (Figure 3A). The increase of *Acod1* expression was next confirmed by RT-qPCR using *Tet2* KO MuSCs (Figure 3B). The corresponding elevation in itaconate level was also detected in *Tet2* KO MuSCs (Figure 3D). When *Acod1* was over-expressed in WT MuSCs via lentivirus infection, succinate level was elevated (Figures 3C and S3B). These results suggest that *Acod1* up-regulation resulted from *Tet2* KO leads to accumulation of succinate.

**Figure 3.**
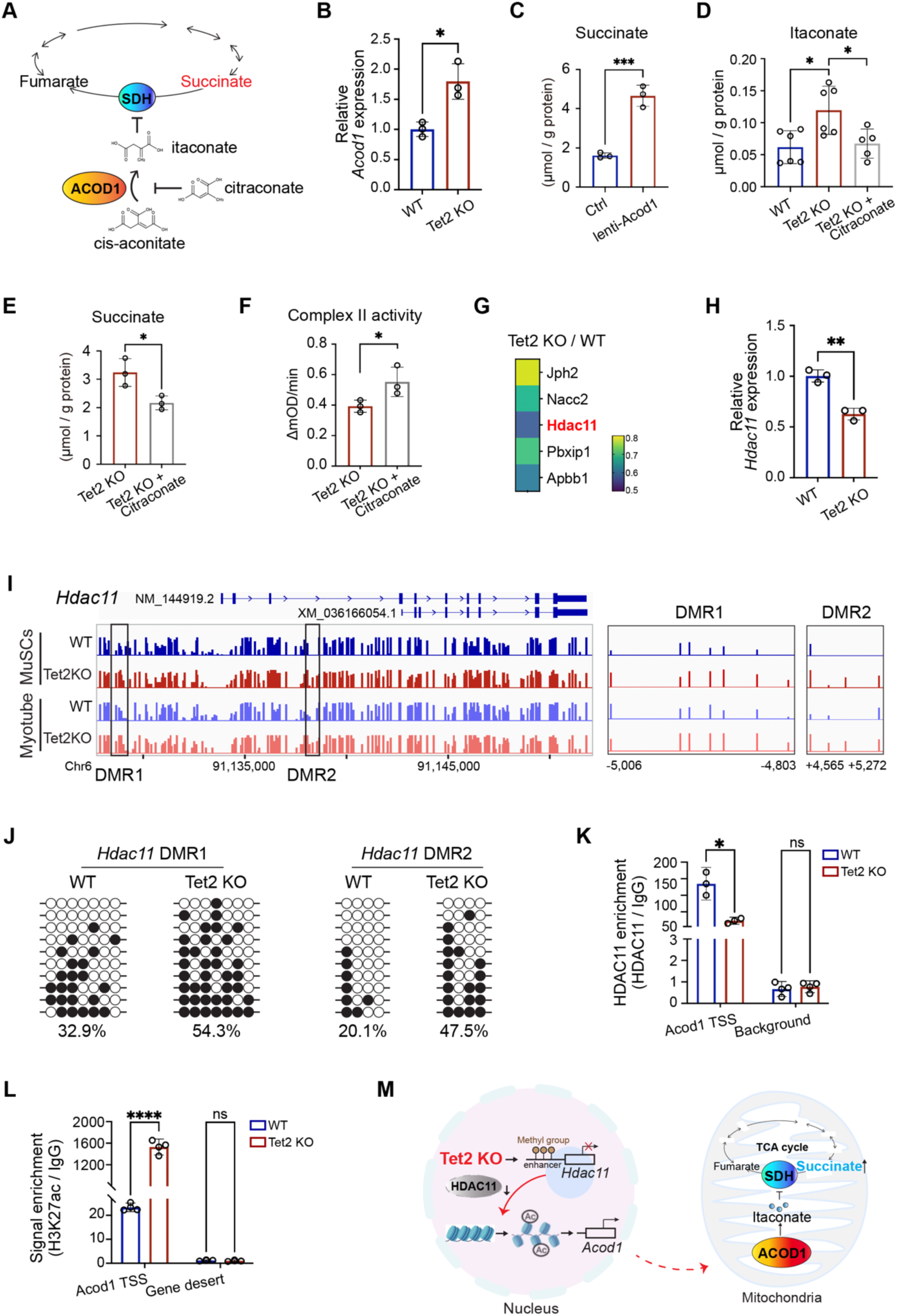
Disruption of a Tet2-Hdac11-Acod1 axis results in succinate accumulation in *Tet2* KO MuSCs. **(A)** Diagram illustrating the role of ACOD1-dependent itaconate production on SDH activity. SDH, succinate dehydrogenase. **(B)** Quantification of relative *Acod1* mRNA level in WT and Tet2 KO MuSCs (n=3 in each group). **(C)** Absolute succinate level measured by coupled enzyme assay in control (Ctrl) and *Acod1* overexpressing MuSCs, respectively (n=3 in each group). MuSCs were infected by lentivirus encoding GFP or *Acod1*. Succinate level was assayed 4 days after virus infection. **(D)** Quantification of targeted LC-MS/MS analysis of itaconate amount in WT, *Tet2* KO, and citraconate treated *Tet2* KO MuSCs (n=6 in each group). **(E)** Absolute succinate content measured by coupled enzyme assay in Tet2 KO and citraconate treated *Tet2* KO MuSCs, respectively (n=3 in each group). **(F)** SDH enzyme activity measured by DCPIP assay in *Tet2* KO and citraconate treated Tet2 KO MuSCs, respectively (n=3 in each group). **(G)** Heatmap analysis of the fold change of gene expression of five hypermethylated transcription repressors in *Tet2* KO compared to that in WT MuSCs. RNA-seq data were from our previously published dataset: GSE158649. **(H)** Analysis of relative *Hdac11* mRNA level in WT and *Tet2* KO MuSCs by RT-qPCR (n=3 in each group). **(I)** Methylation profile of the 5’ upstream region of *Hdac11* gene in WT or Tet2 KO MuSCs and myotubes extracted from genome wide methylation-seq. The left box indicated the methylation profile of *Hdac11* gene in WT and *Tet2* KO MuSCs and myotubes, respectively. The right box indicated the hypermethylated region in *Tet2* KO MuSCs and myotubes highlighted in the black box. And the hypermethylated region was separated into two parts (DMR1 and DMR2). DMR: Differential Methylation Region. **(J)** Bisulfite sequencing results of the DMR1 and DMR2 enhancer of *Hdac11*. White and black circles indicated hypomethylated and hypermethylated CpG sites, respectively. **(K)** Quantification of HDAC11 recruitment on *Acod1* promoter in WT and *Tet2* KO MuSCs by ChIP-qPCR (n=3). TSS, Transcription Start Site. **(L)** Quantification of H3K27ac levels in *Acod1* promoter in WT and *Tet2* KO MuSCs by ChIP-qPCR (n=3). TSS, Transcription Start Site. **(M)** The disrupted enzymatic cascade upon *Tet2* KO ultimately led to abnormal succinate accumulation and mitochondria dysfunction in MuSCs. **(B)-(F), (H), and (K)-(L)** Data were shown in mean ± SD form. Data were analyzed by unpaired t-test. * indicated *p* < 0.05; ** indicated *p* < 0.01; *** indicated *p* < 0.001.

To further prove the notion, *Tet2* KO MuSCs were treated by 1mM citraconate, an Acod1 inhibitor, for 18 hours, and the cell extracts were subjected for subsequent metabolome analysis by mass-spec. PCA analysis indicated that citraconate treatment rescued the global metabolite profile to resemble that of WT MuSCs (Figure S3C and S3D). The levels of itaconate and succinate were down-regulated upon citraconate treatment (Figures 3D, E, and S3D). Consistently, the activity of SDH was also rescued as shown by DCPIP reduction assay (Figure 3F). Taken together, our data suggest that *Acod1* up-regulation results in succinate accumulation in *Tet2* KO MuSCs.

Since Tet2 typically activates transcription of the target genes by active demethylating ^m^C,^1,4^ *Acod1* up-regulation suggests an indirect effect. We therefore searched for transcription repressors down-regulated in our RNA-seq dataset. *Hdac11* (*Histone Deacetylase 11*), which is a transcription inhibitor with histone deacetylase activity, was down-regulated in *Tet2* KO MuSCs (Figures 3G and H). And the enhancer of *Hdac11*, was hypermethylated in *Tet2* KO MuSCs (Figures 3I and J). The *Hdac11* enhancer in WT MuSCs displayed characteristic marks of an active enhancer, with significant enrichment of both H3K4me1 and H3K27Ac (Figures S3E and F). In *Tet2* KO MuSCs, this region displayed a pronounced loss of these active histone marks, consistent with its hypermethylation status (Figures S3G and H). These data suggest that *Hdac11* is the direct target of Tet2.

Hdac11 chromatin immunoprecipitation (ChIP) assays were performed to detect its occupation on *Acod1* promoter. Hdac11 was enriched at the transcription start site (TSS) of *Acod1* in WT MuSCs. In contrast, Tet2 KO led to decreased Hdac11 recruitment at *Acod1* TSS (Figure 3K). The loss of Hdac11 binding led to a marked enrichment of the active histone marks H3K27Ac at *Acod1* TSS (Figure 3L), activating its transcription.

These results delineate an enzymatic cascade that regulates succinate level in MuSCs. Tet2 directly activates the *Hdac11* transcription by removing DNA methylation at its enhancer region. Hdac11 in turn inhibits the transcription of *Acod1*, which is an enzyme catalyzing the production of itaconate, an inhibitor of SDH. Low itaconate level allows SDH efficiently catalyzing succinate dehydrogenation. In *Tet2* KO MuSCs, this enzymatic cascade is disrupted, leading to abnormal succinate accumulation and mitochondria dysfunction (Figure 3M).

### Elevated H4K31succ driven by succinate accumulation activates pro-inflammatory transcription in *Tet2* KO MuSCs

Succinate is a metabolite bridging cell metabolism, epigenetic regulation and cell fate determination.^24,46,47^ Succinate can serve as a substrate to generate histone succinylation, which is dynamically regulated by succinyltransferases CPT1A and KAT2A and erased by desuccinylases SIRT5 and SIRT7 (Figure S4A).^48-52^ The expression of succinyltransferase *Cpt1a* was slightly up-regulated and other related enzymes remained unchanged in *Tet2* KO MuSCs as indicated by RT-qPCR (Figure S4B). As expected, H4K31 succinylation (H4K31succ) level was increased in *Tet2* KO MuSCs as shown by immunoblotting (Figures 4A and B), further confirming the succinate induced H4K31succ elevation in *Tet2* KO MuSCs.

**Figure 4.**
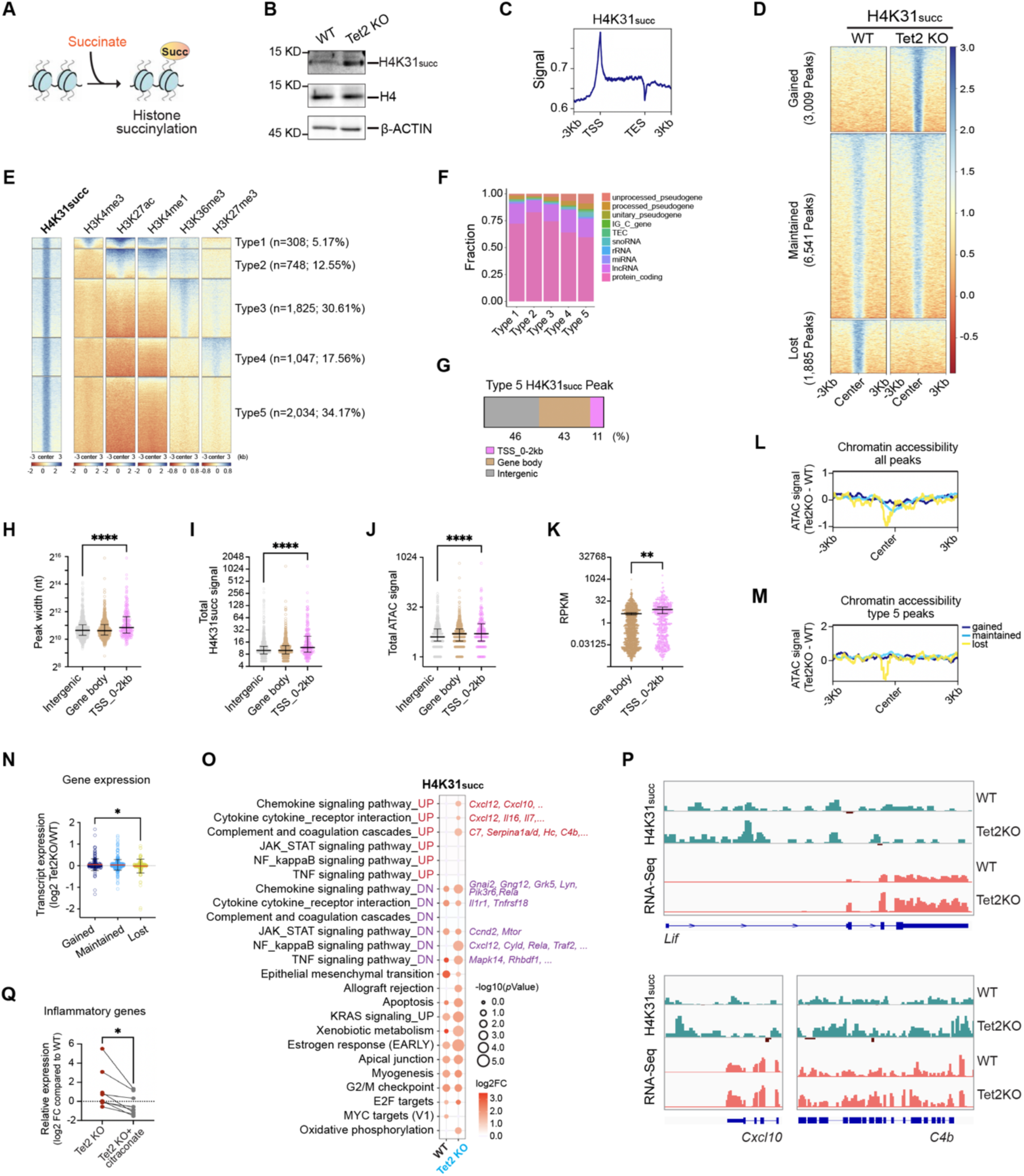
Elevated H4K31succ driven by succinate accumulation activates pro-inflammatory transcription in *Tet2* KO MuSCs. **(A)** Diagram illustrating the histone succinylation process. Red-colored succinate indicated increased succinate level in *Tet2* KO compared to that in WT MuSCs. **(B)** Represented immunoblotting analysis of H4K31succ in WT and *Tet2* KO MuSCs. H4 and β-ACTIN served as loading controls. **(C)** Metagene analysis showing H4K31succ occupancy profiles. Data were from H4K31succ CUT&Tag-seq in WT MuSCs. **(D)** Heatmap representation of H4K31succ enrichment intensity based on CUT&Tag reads in WT and *Tet2* KO MuSCs. Signal within 3kb of the peak center were displayed in descending order for each cluster (i.e., gained, maintained, and lost in response to *Tet2* KO). **(E)** Heatmap representation of 5 types of H4K31succ peaks based on their overlapping with H3K4me3, H3K27ac, H3K4me1, H3K36me3 and H3K27me3 reads. Data were from H4K31succ CUT&Tag-seq in WT MuSCs. **(F)** Percentage of genes by transcript biotype for all annotated genes overlapped with each H4K31succ peak type. **(G)** Genomic feature distribution of type 5 H4K31succ peaks. The relative proportions of type 5 H4K31succ peaks in TSS _0-2Kb (-2000 bp to 0 bp from transcriptional start site), gene body and intergenic regions were displayed. **(H)-(J)** Box plots comparing the peak width (H), H4K31succ CUT&Tag-seq counts (I), or ATAC-seq counts (J) of type 5 H4K31succ peaks in TSS_0-2kb, gene body and intergenic regions. Data were shown in median with interquartile range form. **(K)** Box plots comparing the expression levels of genes with H4K31succ peaks around their transcription start sites (TSS_0-2 Kb) or in their gene bodies (Gene body). Data were shown in mean ± SE form. **(L)-(M)** Metagene analysis of ATAC-seq signal difference between *Tet2* KO and WT MuSCs around all (L) or type 5 (M) H4K31succ peak center in each cluster (i.e., gained, maintained, and lost in response to *Tet2* KO), graphed in 100bp bins. **(N)** Bar plots comparing the expression difference (log2 Tet2 KO/WT) of genes with gained, maintained, and lost H4K31succ peaks (type 5 H4K31succ peaks only) in their TSS or gene body regions upon *Tet2* KO. Data were shown in mean ± SD form. **(O)** Bubble plot comparing functional enrichments for genes with H4K31succ peaks in their TSS or gene body regions in WT and *Tet2* KO MuSCs. Representative genes enriched in each functional term were shown on the right. Dot sizes: -log_10_ (*p* value); dot colors: average log2 fold change (H4K31succ/IgG) of all genes in each functional term (blue to red). **(P)** Genome browser image of H4K31succ CUT&Tag-seq and RNA-seq data at representative inflammatory gene locus in WT and *Tet2* KO MuSCs. **(Q)** Analysis of relative expression of inflammatory genes before and after citraconate treatment by RT-qPCR in *Tet2* KO MuSCs (n=3 in each group). **(H)-(K), (N), and (Q)** Data were analyzed by unpaired t-test. * indicated *p* < 0.05; ** indicated *p* < 0.01; *** indicated *p* < 0.001.

To decode the functional impact of this histone modification genome-wide, H4K31succ CUT&Tag-seq was performed using WT or *Tet2* KO MuSCs. H4K31succ was primarily enriched at TSS (transcription start site) region (Figure 4C). Global increase of H4K31succ peak number was detected in *Tet2* KO MuSCs (Figure 4D), mirroring the immunoblotting data (Figure 4B). Based on the co-localization with canonical histone marks, H4K31succ peaks were categorized to 5 types. Type 1 peaks were co-localized with H3K4me3 and H3K27Ac, associated with active promoters, and accounted for 5.17% of the total peaks. Type 2 peaks were co-localized with H3K27Ac and H3K4me1, associated with active enhancers, and accounted for 12.55% of the total H4K31succ peaks. Type 3 peaks were co-localized with H3K36me3, associated with gene bodies, and accounted for 30.61% of the total peaks. Type 4 peaks were co-localized with H3K27me3, and constituted 17.56% of the total peaks. Type 5 peaks were unique and showed no overlapping with other known histone marks. They accounted for 34.17% of the total H4K31succ peaks (Figures 4E and S4C).

The unique H4K31succ peaks (Type 5) were further analyzed. They were overwhelmingly enriched at protein coding genes, with 11% residing within promoter regions (Figures 4F and G). These peaks exhibited wider peak regions, stronger signals, higher chromatin accessibility, and positive correlation with active transcription of target genes (Figures 4H-K and S4D), suggesting that H4K31succ recruitment on promoters contributes to transcriptional activation. In line with H4K31succ as an active transcription mark, 3,009 gained peaks showed elevated chromatin accessibility in Tet2 KO MuSCs (Figures 4L and M). And a genome-wide positive correlation between the H4K31succ enrichment and transcription activation was observed (Figure 4 N), further supporting that H4K31succ is an active histone mark.

Further analysis of the target genes revealed that H4K31succ was enriched at the promoters of inflammatory genes, such as chemokine, cytokine cytokine_receptor interaction, complement and coagulation cascades, Jak-STAT, NF kappa B, and TNF signaling (Figure 4O), suggesting potential activation of inflammatory genes in *Tet2* KO MuSCs. Consistently, expression of inflammatory genes was up-regulated in these cells (Figure 4P). Citraconate treatment of *Tet2* KO MuSCs reversed the activation of inflammatory genes by reducing cellular succinate level (Figure 4Q). Taken together, these results suggest that accumulation of succinate in *Tet2* KO MuSCs fuels H4K31 succinylation, which in turn activates transcription of pro-inflammatory genes, providing a link between metabolism reprogramming and sterile inflammation.

### Excess succinate from *Tet2* KO MuSCs transforms skeletal muscle into a source of inflammatory signals via cell fusion during homeostasis

The above results demonstrated that excess succinate induced by *Tet2* KO triggered activation of inflammation program in MuSCs (Figure 4). However, given the low number of MuSC *in vivo*,^34,53^ they were unlikely to sustain whole-body chronic inflammation. Thus, we further investigated how the activation of inflammation program in a minor cell population is amplified to exert whole-body effects.

*Tet2* KO was specifically induced in MuSCs of adult mice, leaving pre-existing myofibers which housed large amount of myonuclei untouched. Furthermore, in the absence of injury, only sporadic *Tet2* KO MuSCs fused during routine homeostasis. Consequently, the vast majority of myonuclei should retain WT *Tet2* gene and express normal level of Tet2. To test this, single myofibers from *Tet2* KO mice were isolated and subjected to RT-qPCR examination. As predicted, *Tet2* expression was largely unchanged 3 months after *Tet2* KO in MuSCs (Figure S5A). In sharp contrast, the expression of inflammatory genes was up-regulated in the same myofibers (Figure S5B). These results suggest that the inflammation activation in myofibers is not a consequence of incorporating large number of *Tet2* KO MuSCs and down-regulation of Tet2 as the case in muscle regeneration.

To investigate the alternative mechanism of inflammatory gene activation in myofibers, *Tet2* KO MuSCs were mixed with WT MuSCs at various ratios *in vitro* in differentiation medium to model the sporadic fusion of Tet2 KO MuSCs into WT myofibers *in vivo* (Figure 5A). Strikingly, as little as 1% of *Tet2* KO MuSCs was sufficient to induce a significant shortening of mitochondria length in myotubes, phenocopying the defect observed in *Tet2* KO myotubes (Figures 5B and C). At 10% of *Tet2* KO MuSC incorporation, the defect matched that in *Tet2* KO myotubes (Figures 5B and C). The phenotype cannot be attributed to the loss of *Tet2* in myotubes because *Tet2* expression level was largely unchanged even with up to 10% *Tet2* KO MuSC incorporation (Figure S5C). Expression of inflammatory genes increased in all myotubes containing higher than 1% of *Tet2* KO myonuclei (Figure S5D). Collectively, these results suggest that the mitochondria dysfunction and inflammatory gene activation in myotubes are not driven by reduction of *Tet2* levels, but rather by other non-nucleus cargo transferred from *Tet2* KO MuSC during fusion.

**Figure 5.**
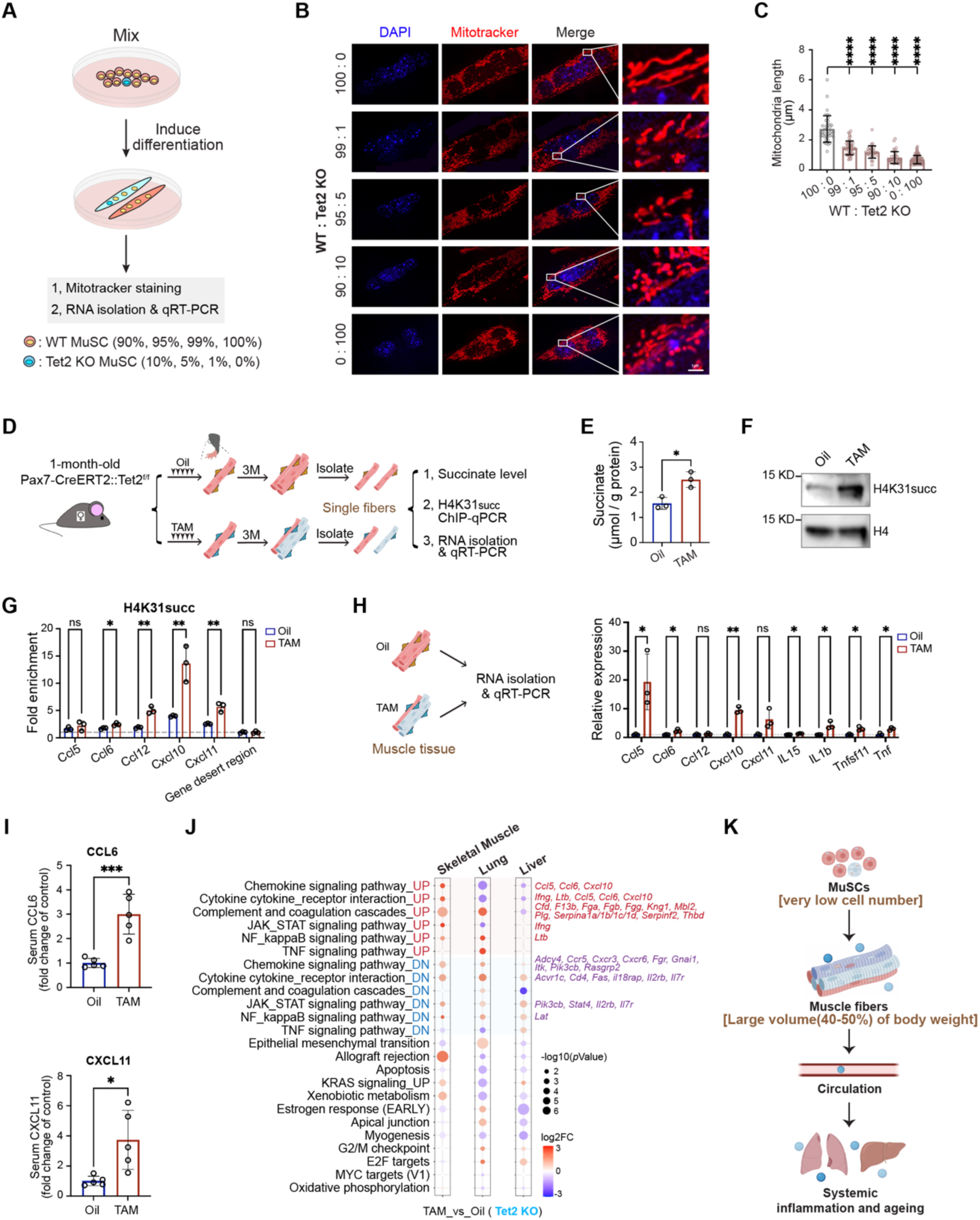
Excess succinate from *Tet2* KO MuSCs transforms skeletal muscle into a source of inflammatory factors via cell fusion during homeostasis. **(A)** Scheme of *in vitro* fusion experiments using variant ratio of Tet2 KO and WT MuSCs. **(B)** Representative images of the mitochondria stained with MitoTracker (red) and DAPI (blue). Scale bars, 1μm. **(C)** Quantification of the mitochondria length. **(D)** Scheme of assays performed in isolated single myofibers from oil or TAM injected mice. **(E)** Absolute succinate content measured by coupled enzyme assays (n=3 in each group). **(F)** Immunoblotting analysis of H4K31succ protein level in isolated single myofibers from oil or TAM injected mice. H4 served as a loading control. **(G)** Quantification of H4K31succ enrichment in TSS regions of *Ccl5*, *Ccl6*, *Ccl12*, *Cxcl10*, and *Cxcl11* by ChIP-qPCR (n=3 in each group). **(H)** Analysis of the relative expression level of inflammatory genes in quadriceps muscle from oil and tamoxifen injected *Pax7* CreERT2:*Tet2* f/f mice (n=3). **(I)** Levels of CCL6 and CXCL11 were analyzed by ELISA using serum from oil or TAM injected *Pax7* CreERT2:*Tet2* f/f mice (n=5). **(J)** Bubble plot comparing functional enrichments for genes differentially expressed (p < 0.05) in quadriceps, lung, and liver between TAM and oil injected Pax7 CreERT2:Tet2 f/f mice. Representative genes enriched in each functional term were shown on the right. Dot sizes: -log_10_ (*p* value); dot colors: average log2 fold change (TAM/Oil) of all genes in each functional term (blue to red). UP: upstream regulators; DN: downstream genes. **(K)** Model graph illustrating the mechanism of inflammation amplification from MuSCs to skeletal muscle. **(C), (E), and (G)-(I)** Data were shown in mean ± SD form. Data were analyzed by unpaired t-test. * indicated *p* < 0.05; ** indicated *p* < 0.01; *** indicated *p* < 0.001.

In cell fusion, not only the nuclei, but also the accumulated succinate and defective mitochondria in *Tet2* KO MuSCs were delivered to myofibers. To test the effects of excess succinate transferred to myofibers, WT myotubes were treated by 0.2mM and 2mM methyl-succinate *in vitro*, respectively. Succinate treatment resulted in shortened mitochondria, recapitulating the defects in *Tet2* KO MuSCs and myotubes (Figures S5E and F). H4K31succ level was elevated (Figure S5G). And the expression of inflammatory genes was subsequently up-regulated after treatment (Figure S5H). These results suggest that excess succinate is sufficient to drive mitochondria defects and activation of inflammatory genes.

This mechanism was further investigated in single myofibers isolated from MuSC specific *Tet2* KO mice (Figure 5D). Succinate level in single myofibers was higher in *Tet2* KO mice compared to that in WT myofibers (Figures 5D and E). And defective mitochondria were widespread in muscle fibers from *Tet2* KO mice (Figure S5I). H4K31succ level increased as indicated by immunoblotting (Figure 5F). The recruitment of H4K31succ to inflammatory genes was enriched in myofibers (Figure 5G). And the expression of inflammatory genes was activated in both single myofibers and the skeletal muscles (Figures 5H and S5B). Collectively, these results support the notion that succinate carried over by *Tet2* KO MuSCs to myofibers during fusion can activate pro-inflammatory program by increasing H4K31succ level.

Via this mechanism, skeletal muscle was converted to be a source of pro-inflammatory factors and increase their serum level as indicated by ELISA assays (Figure 5I). Further RNA-seq analysis from skeletal muscle, lung, and liver from *Tet2* KO mice was performed. The expression of pro-inflammatory factors was elevated in skeletal muscle, while the downstream inflammatory response genes were activated in lung and liver (Figure 5J), causing inflammation and fibrosis in these organs.

Taken together, these results suggest a powerful amplification mechanism mediated by a metabolite. Excess succinate was delivered to myofibers by sporadic *Tet2* KO MuSC fusion during muscle homeostasis. Excess succinate then drove mitochondria defects and H4K31succ mediated epigenetic reprogramming, and transformed skeletal muscle into a potent source of pro-inflammatory signals, thereby propagating and sustaining inflammaging throughout the body (Figure 5K).

### Citraconate treatment rescues systemic inflammaging and premature aging phenotypes in MuSC specific *Tet2* KO mice

The abnormal accumulation of succinate and impaired SDH activity in skeletal muscles were further examined by seahorse assays and metabolome analysis using muscle from MuSC specific *Tet2* KO mice (Figure S6A). SDH defects and succinate accumulation were detected in these mice (Figures S6B-E). Using this mouse model, we further investigated whether the premature aging phenotype could be rescued by increasing SDH activity with citraconate. Intraperitoneally (ip) injection of 50mg/kg body weight of citraconate was administrated to *Pax7* CreERT2: *Tet2* f/f mice once every week for 8 times after TAM induction of *Tet2* KO in MuSCs (Figure 6A). Succinate level in muscles was efficiently reduced in *Tet2* KO mice after citraconate injection (Figure 6B). Consequently, the levels of H4K31succ and expression of inflammatory genes were also down-regulated (Figures 6C and D). Notably, the premature aging phenotypes including body weight reduction, hair loss and the development of hunched back in MuSC specific *Tet2* KO mice were all rescued by citraconate injection (Figures 6E-H). Furthermore, functional and structural metrics of muscle health, such as running distance, running time, muscle weight, and muscle fiber size were all recovered after citraconate injection (Figures 6I-K). Collectively, these data suggest that citraconate injection rescues the premature aging phenotypes.

**Figure 6.**
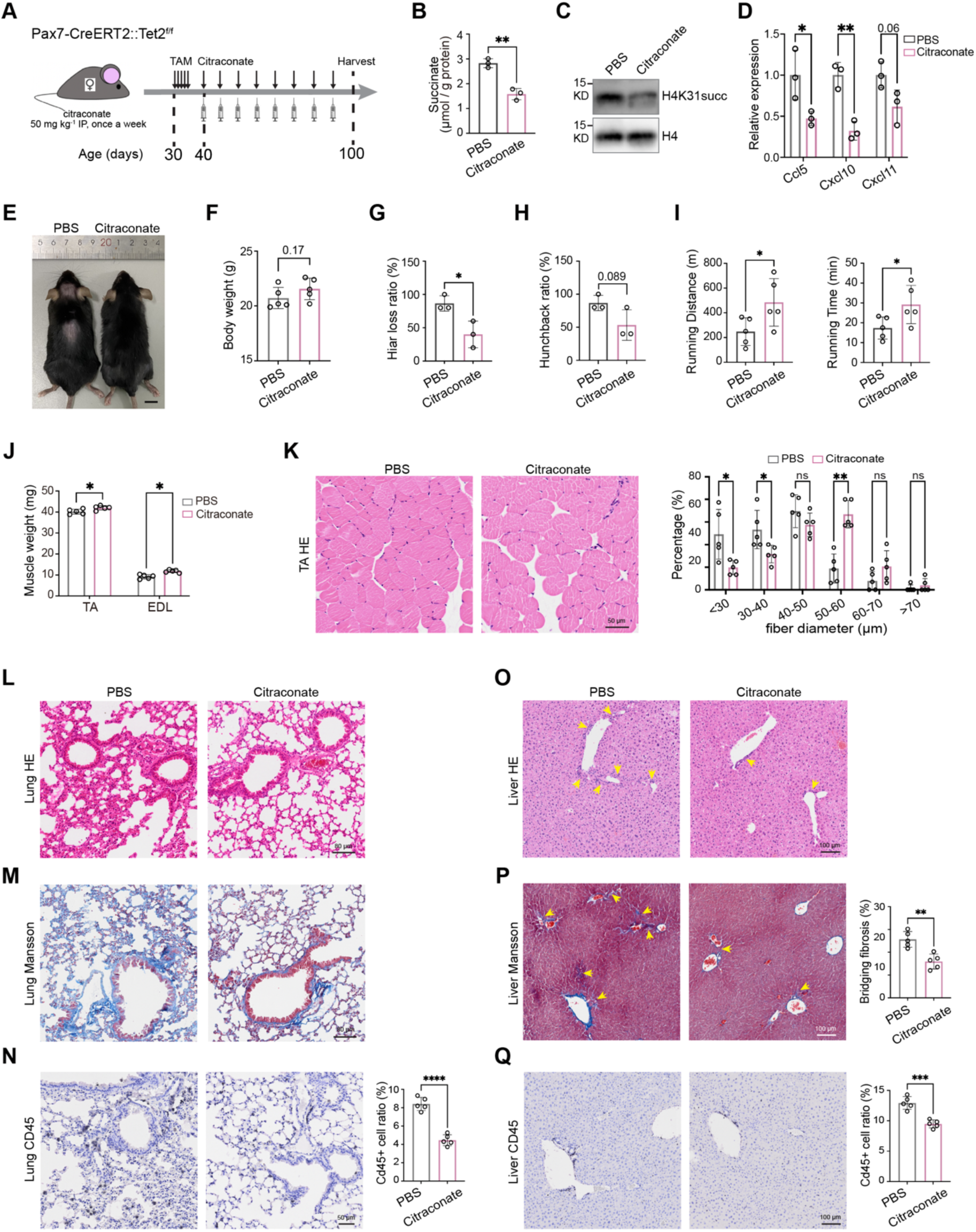
Citraconate treatment rescues systemic inflammaging and premature aging phenotypes in MuSC-specific *Tet2* KO mice. **(A)** Scheme of citraconate treatment experiment. Female Pax7 CreERT2:Tet2 f/f mice were injected with TAM at the age of 1 month (30-day-old). Then, mice were given either PBS or citraconate (50 mg/kg) starting from 5 days after the last TAM injection (40-day-old) once a week for 8 times. Mice were harvested 11 days after the last injection (100-day-old). **(B)** Absolute succinate content measured by coupled enzyme assays in quadriceps with PBS or citraconate injection in Tet2 KO mice, respectively (n=3 in each group). **(C)** Immunoblotting of H4K31succ protein in quadriceps from Tet2 KO mice with PBS or citraconate injection. H4 served as a loading control. **(D)** Analysis of the relative expression of *Ccl5*, *Cxcl10*, and *Cxcl11* gene by RT-qPCR in quadriceps from Tet2 KO mice with PBS or citraconate injection (n=3 in each group). **(E)** Representative images of 100-day-old MuSC specific Tet2 KO mice with PBS or citraconate injection. Scale bar: 1cm. **(F)** Statistical analysis of body weights of Tet2 KO mice with PBS or citraconate injection (n = 5 in each group). p=0.17 using unpaired t-test analysis. **(G)** Statistical analysis of hair loss ratio of Tet2 KO mice with PBS or citraconate injection. The ratio was calculated based on data collected from 3 batches of mice, 5 mice in each batch. **(H)** Statistical analysis of hunchback ratio of Tet2 KO mice with PBS or citraconate injection. The ratio was calculated based on data collected from 3 batches of mice, 5 mice in each batch. p=0.089 using unpaired t-test analysis. **(I)** Mean running time and running distance of Tet2 KO mice with PBS or citraconate injection on a motorized treadmill (n = 5 in each group). **(J)** Extensor digitorum longus (EDL) and TA muscle weights of Tet2 KO mice with PBS or citraconate injection (n = 5 in each group). **(K)** Representative images of H&E staining of TA muscles Tet2 KO mice with PBS or citraconate injection (left), and percentage quantification of myofiber size (right) (n=5). Scale bar, 50μm. **(L)-(M)** Representative images of H&E (L) and Masson (M) staining of lung from 100-day-old Tet2 KO mice with PBS or citraconate injection. Scale bars, 80μm. **(N)** Representative images of CD45 immunohistochemistry staining in lung from100-day-old Tet2 KO mice with PBS or citraconate injection (left), and the quantification of the percentage of CD45-positive cells (right). n=5 in each group. Scale bar, 50μm. **(O)** Representative images of H&E staining of liver from 100-day-old Tet2 KO mice with PBS or citraconate injection. For H&E staining, area with potential immune cell infiltration was indicated by yellow arrows. Scale bar, 100μm. **(P)** Representative images of Masson staining of liver from 100-day-old Tet2 KO mice with PBS or citraconate injection were shown at the left. Portal triads to Central vein (P-C) bridging fibrosis, which is a marker for severe liver fibrosis, was indicated by yellow arrows. Quantification of the percentage of bridging fibrosis was shown at the right (n=5 in each group). Scale bars, 100μm. **(Q)** Representative images of CD45 immunohistochemistry staining in liver from 100-day-old Tet2 KO mice (left), and the quantification of the percentage of CD45-positive cells (right). n=5 in each group. Scale bars, 100μm. **(B), (D), (G), (I)-(K), (N), and (P)-(Q)** Data were shown in mean ± SD form. Data were analyzed by unpaired t-test. * indicated *p* < 0.05; ** indicated *p* < 0.01; *** indicated *p* < 0.001.

The multi-organ benefits of citraconate treatment were also examined. H&E staining, Masson staining, and CD45 immunohistological staining illustrated that the infiltration of immune cells and fibrosis in both lung and liver in *Tet2* KO mice were significantly reduced following citraconate treatment (Figures 6L-Q).

Taken together, these results suggest that citraconate mediated inhibition of itaconate-succinate axis attenuates systemic inflammaging and alleviates the multi-organ fibrosis and premature aging.

### Age associated decline of *Tet2* in MuSCs induces multi-organ inflammaging during aging

The premature aging observed in MuSC specific *Tet2* KO mice suggests that down-regulation of *Tet2* in MuSCs could play a role in natural aging. We therefore set out to investigate whether the same pathway is activated during natural aging. The expression of *Tet2* was evaluated by RT-qPCR using MuSCs isolated from young or old murine skeletal muscle, respectively (Figure 7A). *Tet2* expression was reduced sharply in old MuSCs (Figure 7B). In the same line of mechanism, the age associated *Tet2* expression decline was accompanied by *Hdac11* down-regulation and *Acod1* up-regulation (Figure 7B). Consequently, succinate and the subsequent H4K31succ levels surged in old MuSCs (Figures 7C and D). H4K31succ cut-tag sequencing analysis was next performed using MuSCs isolated from young or old mice, respectively. The enrichment of H4K31succ at the promoters of inflammatory genes was detected (Figure 7E), recapitulating the pattern observed in *Tet2* KO MuSCs. Consistent with the epigenetic alteration, the expression of inflammatory genes was up-regulated in old MuSCs (Figure 5F). Analysis of an RNA-seq dataset in public database (accession number GSE183643)^54^ further confirmed the activation of pro-inflammatory genes in old MuSCs, mirroring the transcription signature of *Tet2* KO MuSCs (Figure 7G).

**Figure 7.**
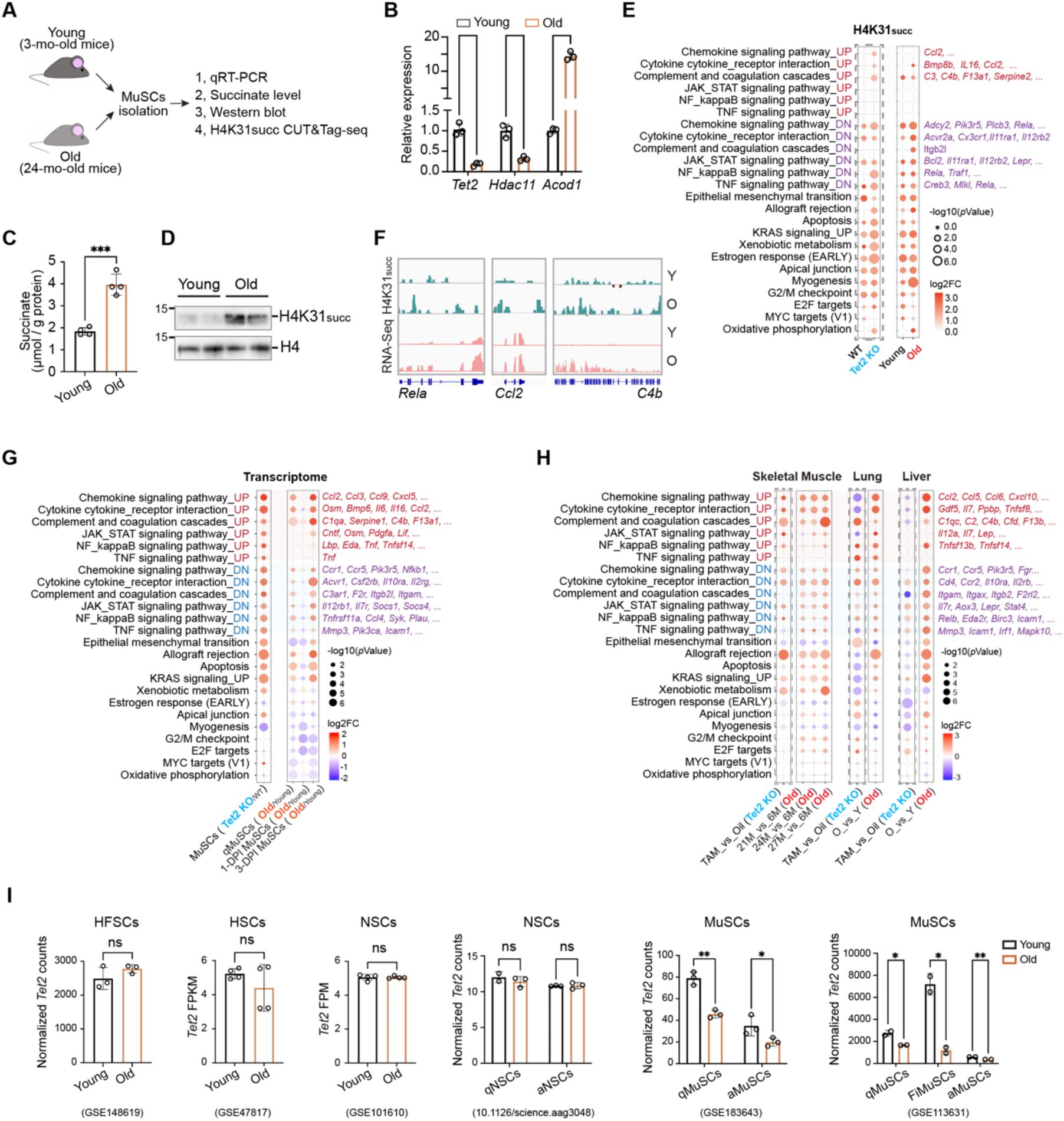
Age-associated decline of *Tet2* expression in MuSCs induces multi-organ inflammaging during aging. **(A)** Scheme of experiments. **(B)** Analysis of the relative expression level of *Tet2*, *Hdac11*, and *Acod1* in young and old MuSCs, respectively (n=3 in each group). **(C)** Absolute succinate content measured by coupled enzyme assays in young and old MuSCs, respectively (n=4 in each group). **(D)** Immunoblotting analysis of H4K31succ in young and old MuSCs. H4 served as a loading control. **(E)** Bubble plot comparing functional enrichments for genes with H4K31succ peaks in their TSS or gene body regions in young and old MuSCs. Representative genes enriched in each functional term were shown at the right. As a control, functional enrichments for H4K31succ peaks in WT and *Tet2* KO MuSCs were marked by grey dashed box. Dot sizes: -log_10_ (*p* value); dot colors: average log2 fold change (H4K31succ/IgG) of all genes in each functional term (blue to red). **(F)** Genome browser image of H4K31succ CUT&Tag-seq and RNA-seq data at representative inflammatory gene locus in young and old MuSCs. **(G)** Bubble plot comparing functional enrichments for genes differentially expressed (p<0.05) between WT and *Tet2* KO MuSCs, young and old qMuSCs, 1-DPI-MuSCs, 3-DPI-MuSCs. Representative genes enriched in each functional term were shown at the right. Dot sizes: -log_10_ (*p* value); dot colors: average log2 fold change (*Tet2* KO/WT or Old/Young) of all genes in each functional term (blue to red). RNA-seq data of WT and *Tet2* KO MuSCs, young and old MuSCs were from GSE158649 and GSE183643, respectively. qMuSCs indicated quiescent MuSCs; 1-DPI-MuSCs indicated MuSCs isolated from mice 1 day post injury; 3-DPI-MuSCs indicated MuSCs isolated from mice 3 days post injury. **(H)** Bubble plot comparing functional enrichments for genes differentially expressed (p<0.05) in skeletal muscle, lung, and liver between young and old mice. Representative genes enriched in each functional term were shown at the right side. Functional enrichments for genes differentially expressed (p<0.05) between oil and TAM injected Pax7 CreERT2:Tet2 f/f mice for each tissue were marked by grey dashed box. Dot sizes: -log_10_ (*p* value); dot colors: average log2 fold change (Old/Young) of all genes in each functional term (blue to red). RNA-seq data of young and old skeletal muscle, lung, and liver were from GSE226117, GSE124872, and PRJNA281127, respectively. **(I)** Relative mRNA level of *Tet2* in HFSCs, HSCs, NSCs, and MuSCs isolated from young and old mice, respectively. All data were from published datasets. The accession numbers were listed below each bar graph. qNSCs indicated quiescent NSCs; aNSCs indicated activated NSCs; qMuSCs indicated quiescent MuSCs; aMuSCs indicated activated MuSCs; FiMuSCs indicated freshly isolated MuSCs. **(B), (C), and (I)** Data were shown in mean ± SD form. Data were analyzed by unpaired t-test. * indicated *p* < 0.05; ** indicated *p* < 0.01; *** indicated *p* < 0.001.

Similar to the scenario in the muscles from MuSC specific *Tet2* KO mice, the expression levels of *Tet2*, *Hdac11*, and *Acod1* were largely unchanged in muscle fibers isolated from old mice (Figure S7A), whereas succinate level was elevated significantly in old skeletal muscles (Figure S7B). The RNA-seq data of organs from old mice were analyzed. The expression of pro-inflammatory factors was up-regulated in muscle (Figure 7H). The downstream inflammatory response genes were activated in lung and liver (Figure 7H). Collectively, these data suggest that inflammation gene activation driven by Tet2-succinate axis is engaged during physiological aging.

We next investigated whether *Tet2* down-regulation is a general feature of aging across adult stem cell types. The expression level of *Tet2* in hair follicle stem cells (HFSCs),^55^ hematopoietic stem cells (HSCs),^56^ neural stem cells (NSCs),^57,58^ and MuSCs from young or old mice^54,59^ was surveyed using available RNA-seq data in the database. Tet2 was specifically down-regulated in old MuSCs (Figure 7I). *Tet2* level in other types of adult stem cells remained to be unchanged during aging (Figure 7I). These results suggest that Tet2-succinate-inflammaging axis is a distinct mechanism in MuSCs. Through the cell fusion in muscle homeostasis, the metabolism disorder driven by *Tet2* down-regulation in aged MuSCs was delivered to muscle, converting skeletal muscle to be a potent source of inflammatory factor production driving whole-body inflammaging and aging.

We further investigated whether the similar mechanism applying to human aging. *TET2* expression level was examined by RT-qPCR using MuSCs isolated from young or old persons, respectively. The expression of *TET2* decreased significantly in MuSCs isolated from old human skeletal muscle compared to that in young MuSCs (Figure S7C). The decline of *TET2* level was associated with increased H4K31succ level and elevated expression of inflammatory genes in old muscles (Figures S7 D and E). These data suggest that the inflammation amplification mechanism originated from *TET2* down-regulation in MuSCs is likely also functioning in human aging.

## DISCUSSION

Inflammaging (sustained chronic low-grade inflammation) is a hallmark and a driving force of aging, contributing to age related multi-organ dysfunction.^10-12^ However, the mechanism that initiates and sustains the sterile inflammatory state remains poorly understood. Here, we uncovered a mechanism that amplifies inflammatory signals originated from a small subset of MuSCs exhibiting age-related *Tet2* downregulation. In these aged MuSCs, aberrant succinate accumulation driven by dysregulation of Tet2-Hdac11-Acod1-SDH axis of enzymatic reactions increased H4K31succ level, remodeling the epigenetic landscape and activating inflammatory gene expression. During muscle homeostasis, these aged MuSCs acted as Trojan horses that delivered the succinate cargo to myofibers upon cell fusion. The metabolite transfer initiated a self-reinforcing cycle of succinate dependent epigenetic reprogramming and inflammatory gene activation in myofibers, thereby transforming skeletal muscle tissue to a potent and persistent source of pro-inflammatory factors. This mechanism fuels systemic inflammation, accelerates multi-organ fibrosis and aging (Figure S7F).

During aging, progressive decline in adult stem cell function, including altered quiescence, self-renewal, and differentiation capacity, leads to impaired tissue regeneration.^60^ Similarly, MuSCs exhibit a sharp functional decline with advanced age, accompanied by increased inflammatory gene expression.^30^ Like most of the adult stem cells, MuSCs account a minor cell population in an organism. Their inflammatory gene activation in MuSC has been viewed primarily as a driver of declined regeneration capacity. The impact of MuSCs on systemic inflammation is negligible.^61,62^ The muscle regeneration decline due to the dysfunction of MuSCs would not be expected to drastically influence aging process under the non-injury condition. Our findings, however, reveal a paradigm shifting role of MuSCs that they exploit the physiological process of cell fusion to broadcast the metabolic stress in a small subset of MuSCs to the entire muscle tissue and promote sustained inflammation. Excess succinate serves as a messenger coupling epigenetic remodeling in a few aged MuSCs to a persistent inflammatory response in a bulk tissue.

MuSC fusion with pre-existing myofibers occurs constitutively in adult muscle during homeostasis without injury. It has been shown that from 6 to 12 months of age, MuSCs contribute to about 25%, 30%, 32% and 19% of myofibers in tibialis anterior (TA), extensor digitorum longus (EDL), diaphragm, and extraocular (EOM) in mice, respectively. This contribution escalates to 75% in the diaphragm by 20 months of age.^63^ This continuous fusion throughout life provides a route through which metabolic alterations in the rare MuSC population are transferred to the myofibers. Given the massive volume and secretory capacity of skeletal muscle, aged MuSCs hijack this homeostatic mechanism to dramatically amplify inflammatory signaling. Thus, through cell fusion, age-related metabolic perturbations in MuSCs, a minor cell population, are directly transmitted to myofibers, positioning MuSCs and skeletal muscle as central regulators of organism inflammation and aging.

Succinate is increasingly recognized as a signaling metabolite mediating mitochondria-nuclear communication.^64^ Our data revealed a pathogenic feedback loop in which excess succinate induced mitochondria dysfunction, which in turn exacerbates succinate accumulation. Targeting this circuit may help develop strategies for aging intervention. As a catalytic product of α-KG-dependent dioxygenases, succinate inhibits histone demethylases and TET family of 5-methlycytosine (5mC) hydroxylases.^23,65^ Succinate also serves as substrate for protein succinylation. Protein succinylation participates in regulation of many cellular processes such as proliferation, differentiation, inflammation, and tumorigenesis.^25-27^ Structure analysis predicts that histone lysine succinylation brings in a negative charge and a bulky side chain to promote DNA unwrapping.^66-68^ Consistent with this prediction, we found that H4K31succ is an active histone mark and a key trigger for inflammatory gene activation in muscle cells. These findings provide fundamental new insights into how metabolite coordinated epigenetic regulation governs tissue homeostasis and aging.

This study suggests that preserving MuSC function is critical not only for maintaining muscle regenerative capacity but also for preventing systemic aging. These findings position MuSC as a key regulator of organism inflammation. Interventions aimed at improving MuSC functions through metabolism remodeling or cell therapy have potentials to alleviate systemic inflammation and decelerate the aging process.

### Limitations of the study

In this study, we focused on the functions of succinate and the consequent epigenetic reprogramming during aging. Mitochondria defects were induced by excess succinate, which in turn could exacerbate metabolic disturbances in aged MuSCs. In this self-reinforcing cycle, besides enhanced succinate accumulation, other metabolic dysfunction also existed. They could collectively accelerate aging. In the future, to further elucidate the link between metabolic shifts and aging will deepen our understanding on the onset and progression of aging and reveal new targets for potential rejuvenation strategies.

## RESOURCE AVAILABILITY

### Lead contact

Further information and requests for resources and reagents should be directed to and will be fulfilled by the lead contact, Ping Hu.

### Materials availability

This study did not generate new, unique reagents.

### Data and code availability

- The H4K31succ CUT&Tag-seq and tissue RNA-seq data from this publication have been deposited into the Genome Sequence Archive in National Genomics Data Center of China under the identifier GSA: CRA034197.
- This paper does not report original code.
- Any additional information required to reanalyze the data reported in this paper is available from the lead contact upon request.

## ACKNOWLEDGMENTS

We thank Drs. Xingguo Liu from Guangzhou Institute of Biomedicine and Health, Chinese Academy of Sciences; Yunshen Chan from Guangzhou National Laboratory for helpful discussions; cell biology facility of Guangzhou National Laboratory for helps with MuSCs sorting; molecular biology facility of Shanghai Institute of Biochemistry and Cell Biology for helps on transmission electron microscopy analysis; molecular biology facility of Guangzhou National Laboratory for helps on seahorse analysis. We thank BioRender (BioRender.com) for providing elements used to create figures in this article. This work is supported by the R&D program of Guangzhou National Laboratory (GZNL2023A02004), the National Natural Science Foundation of China (32170904 and 82361148131).

## AUTHOR CONTRIBUTIONS

Conceptualization, P.H.; Investigation, X.-F.W., M. C., R.-Q. F., H.-Y.W., R.-M.M., H.W., W.W., J.-M.W., G.-L.X., and P.H.; Data analysis, X.-F.W., M.C., and P.H.; Writing – Original Draft, P.H. and X.-F.W.; Writing – Review & Editing, P.H. and X.-F.W.; Supervision, P.H.; Funding Acquisition, P.H.

## STAR★METHODS

### KEY RESOURCES TABLE

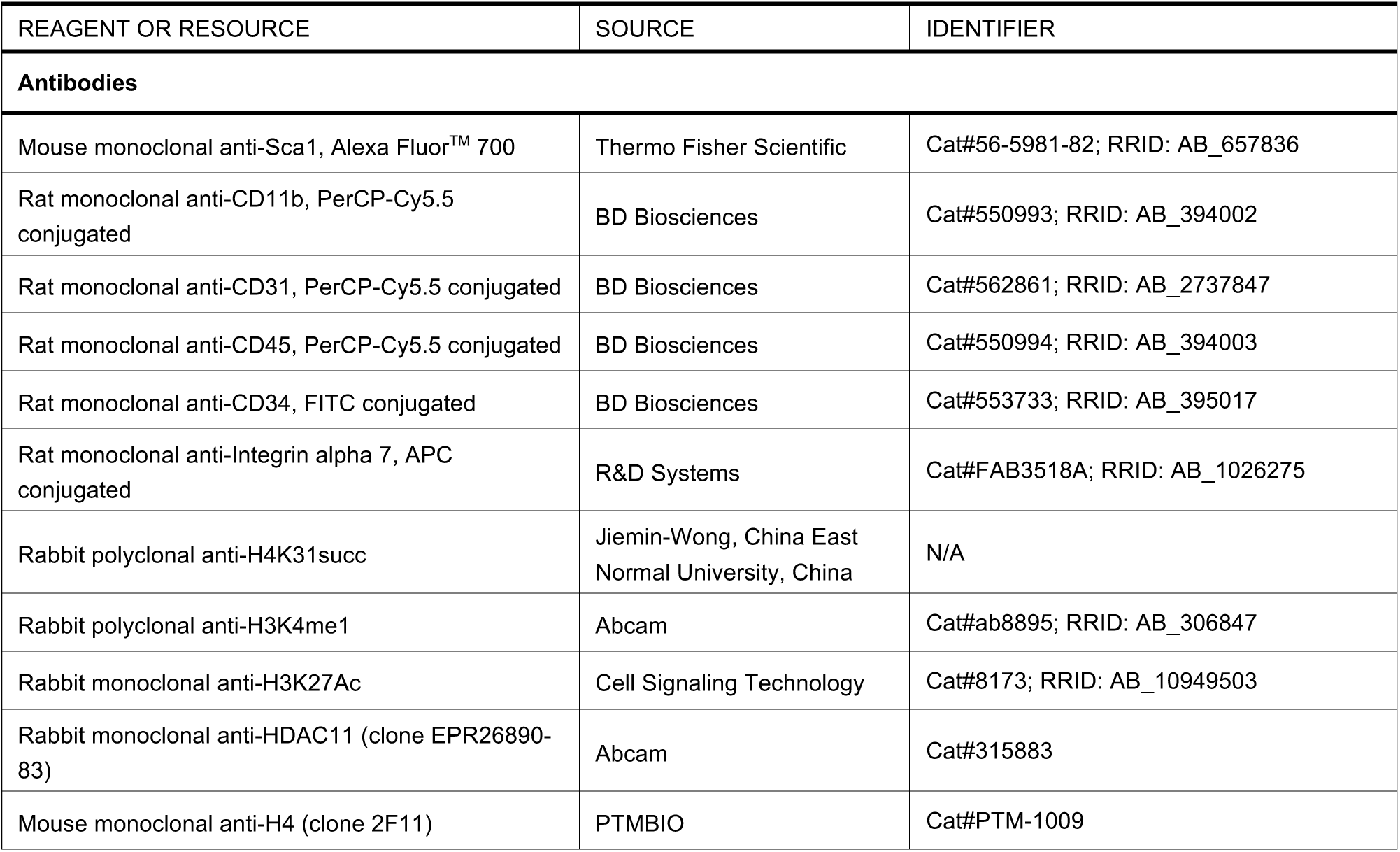
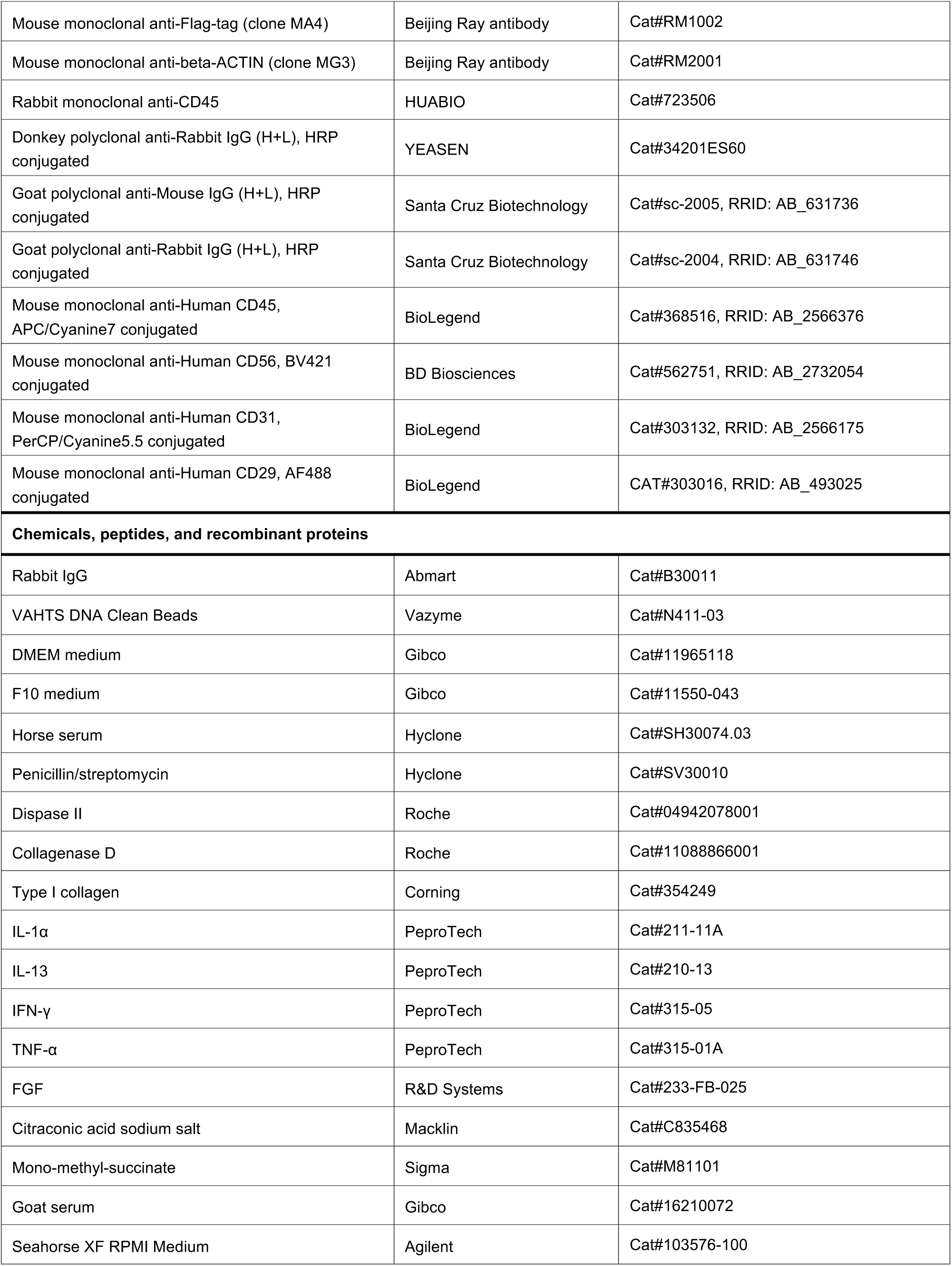
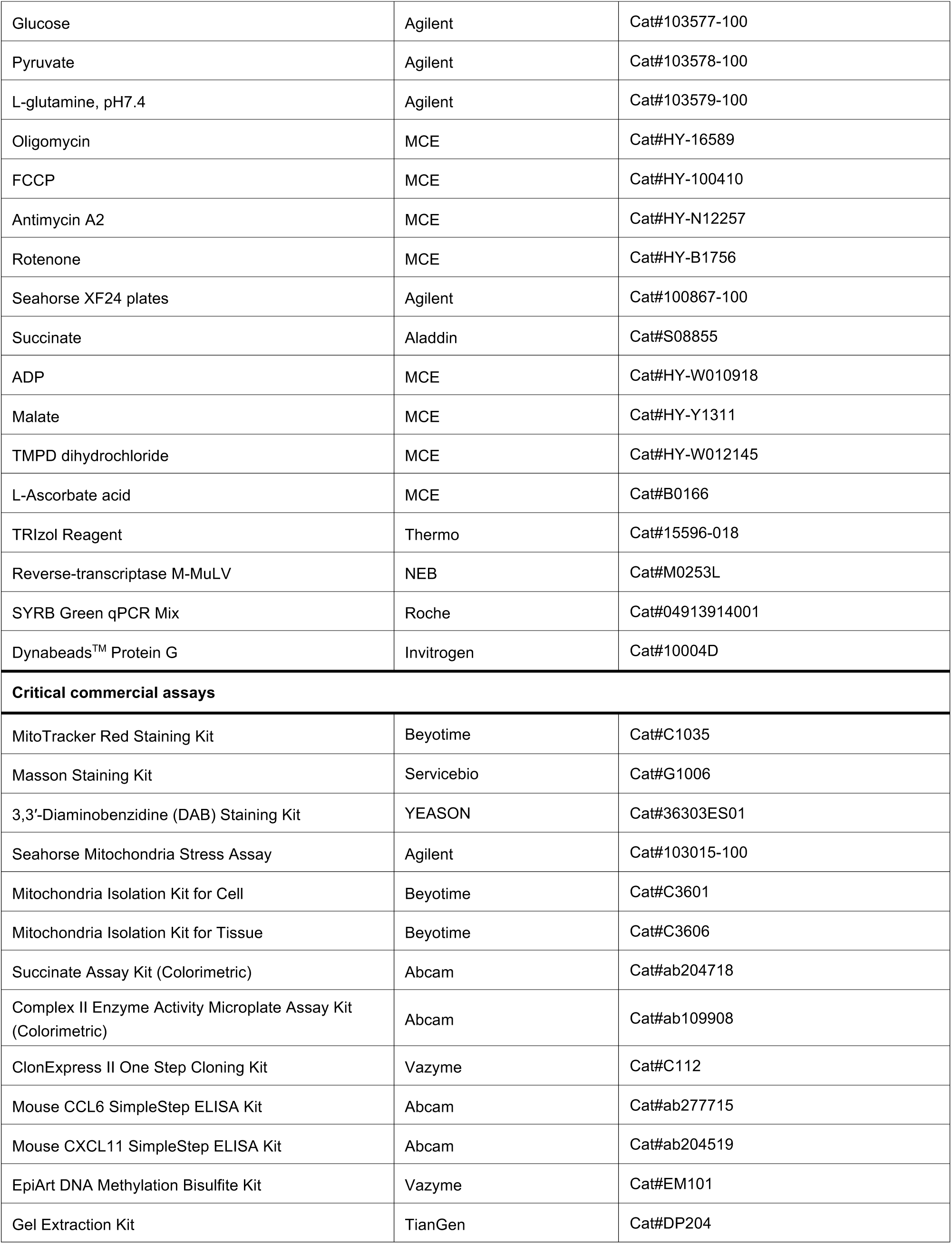
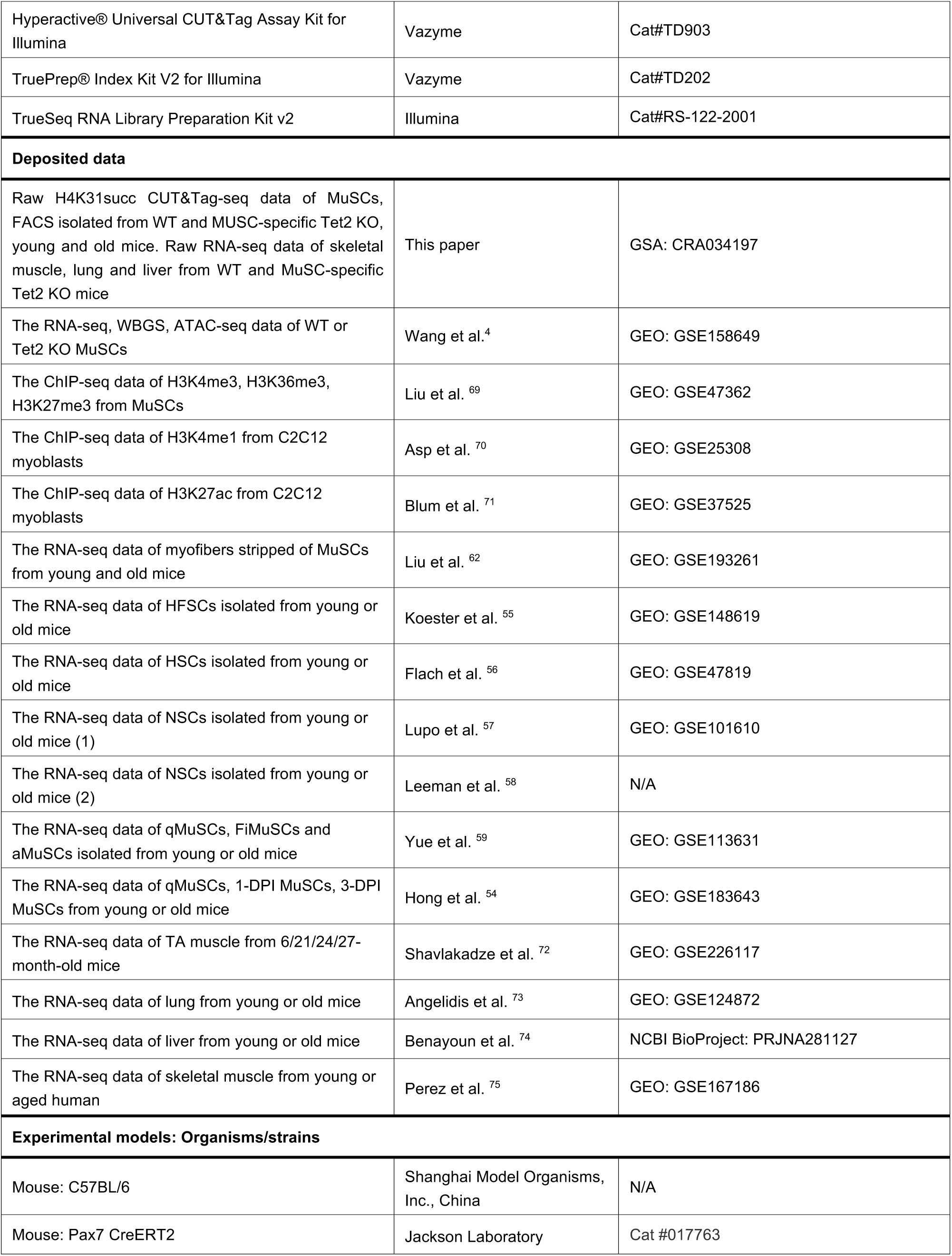
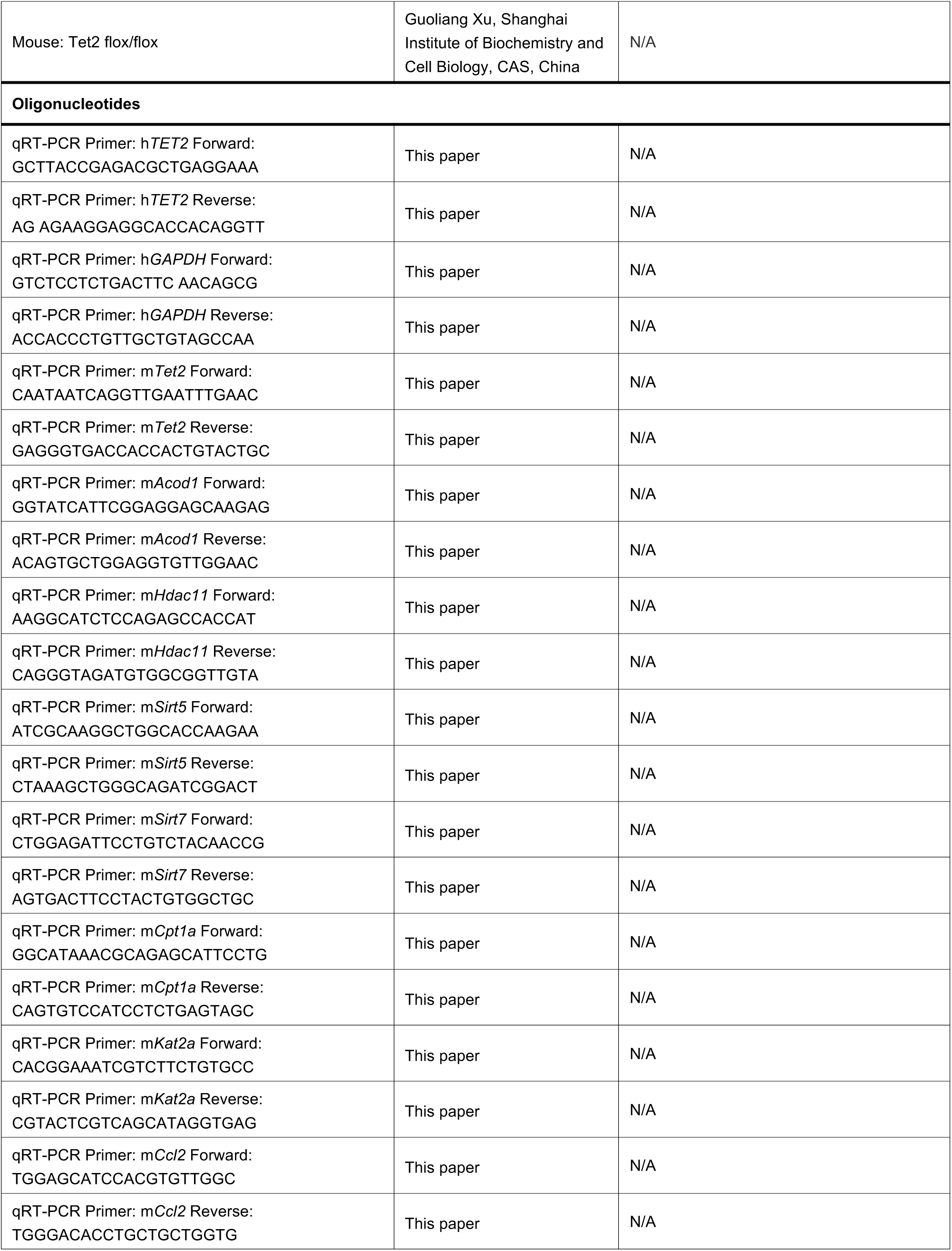
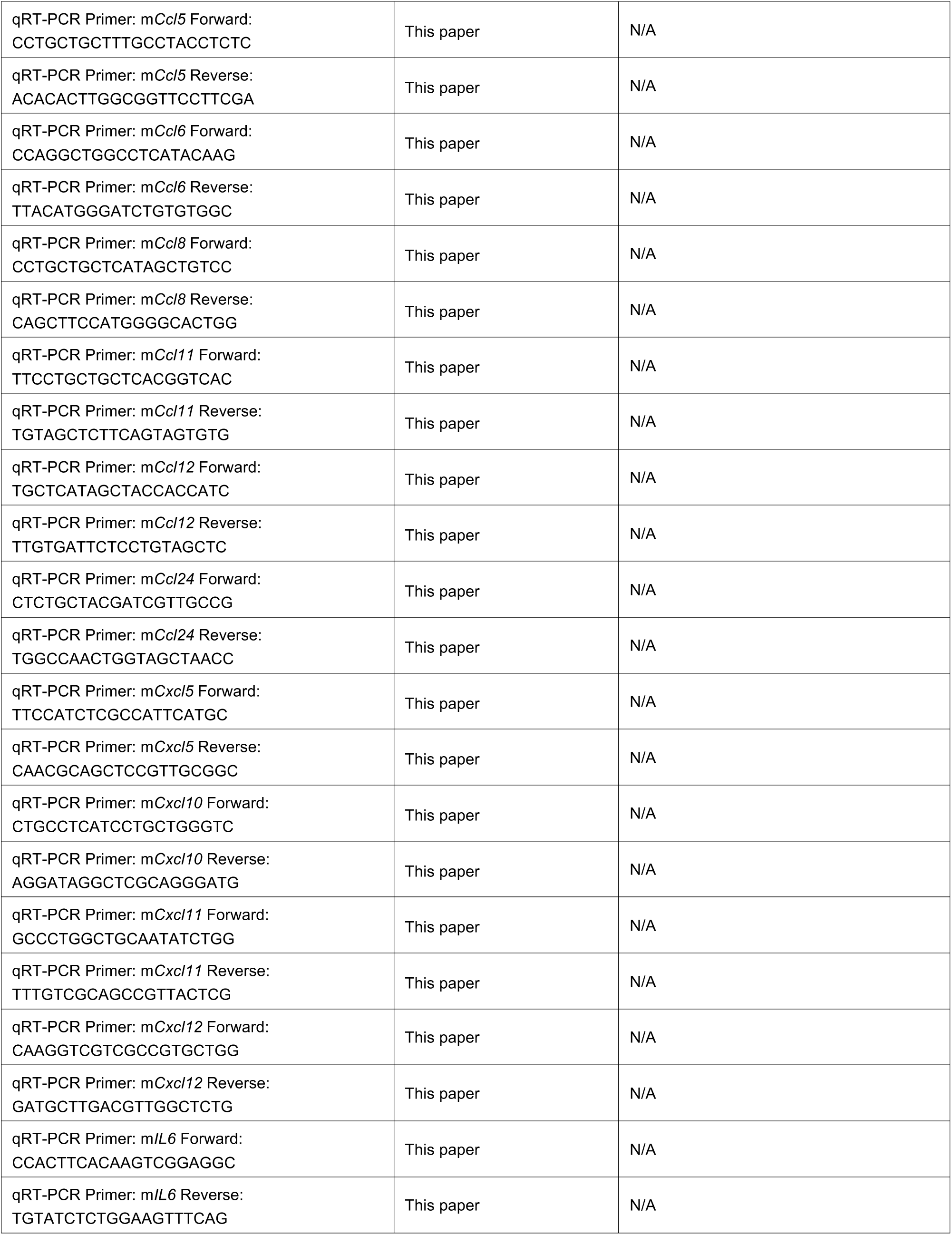
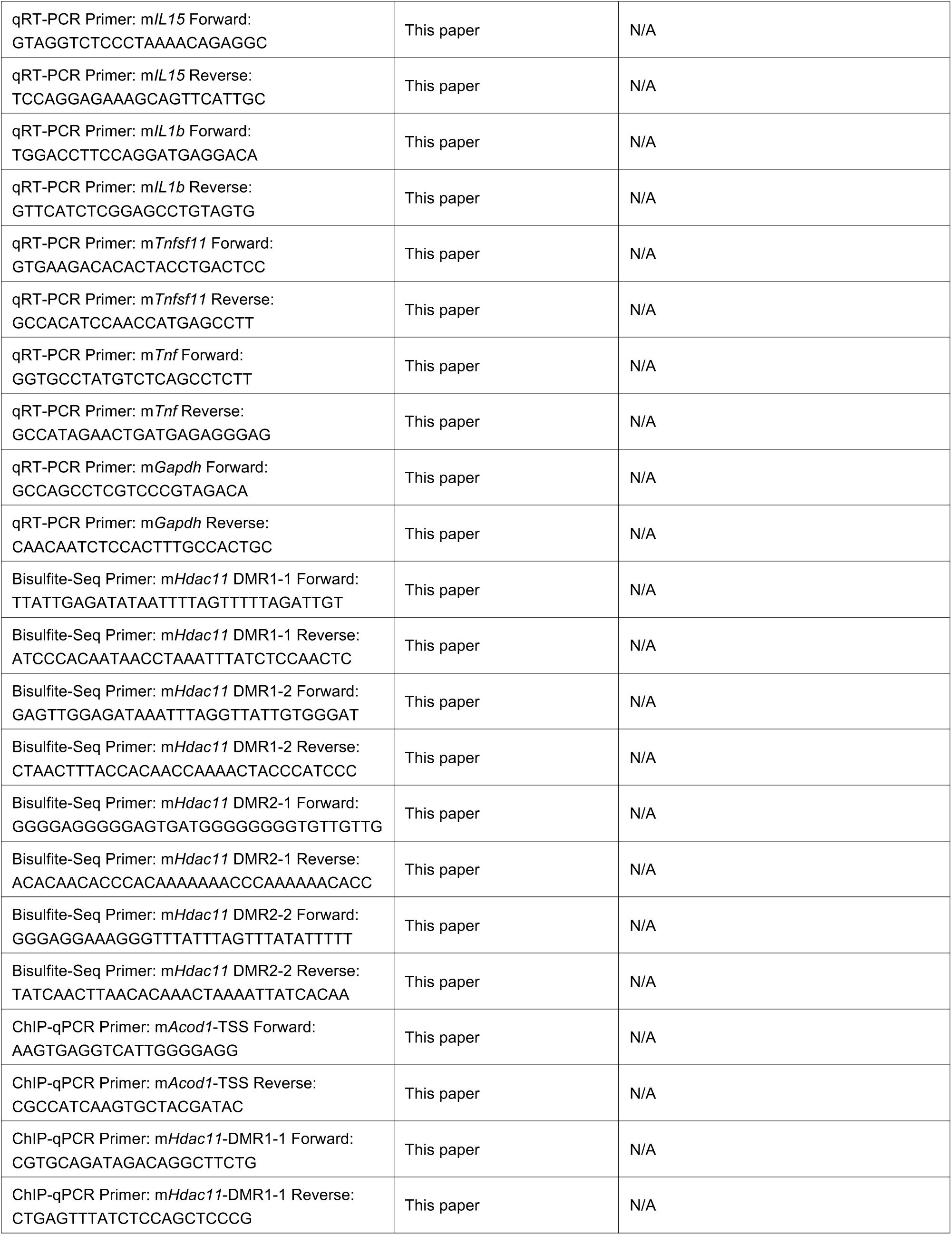
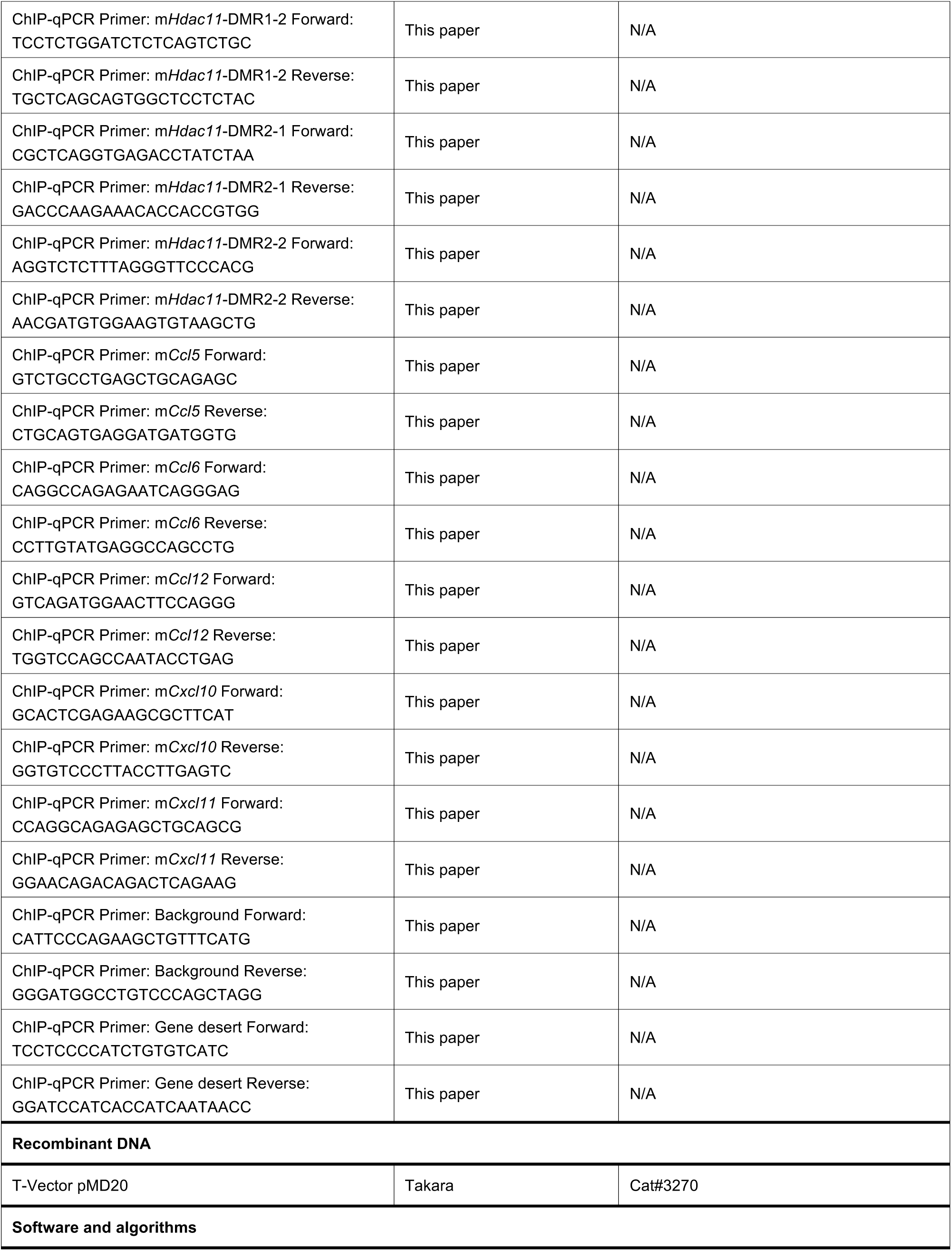
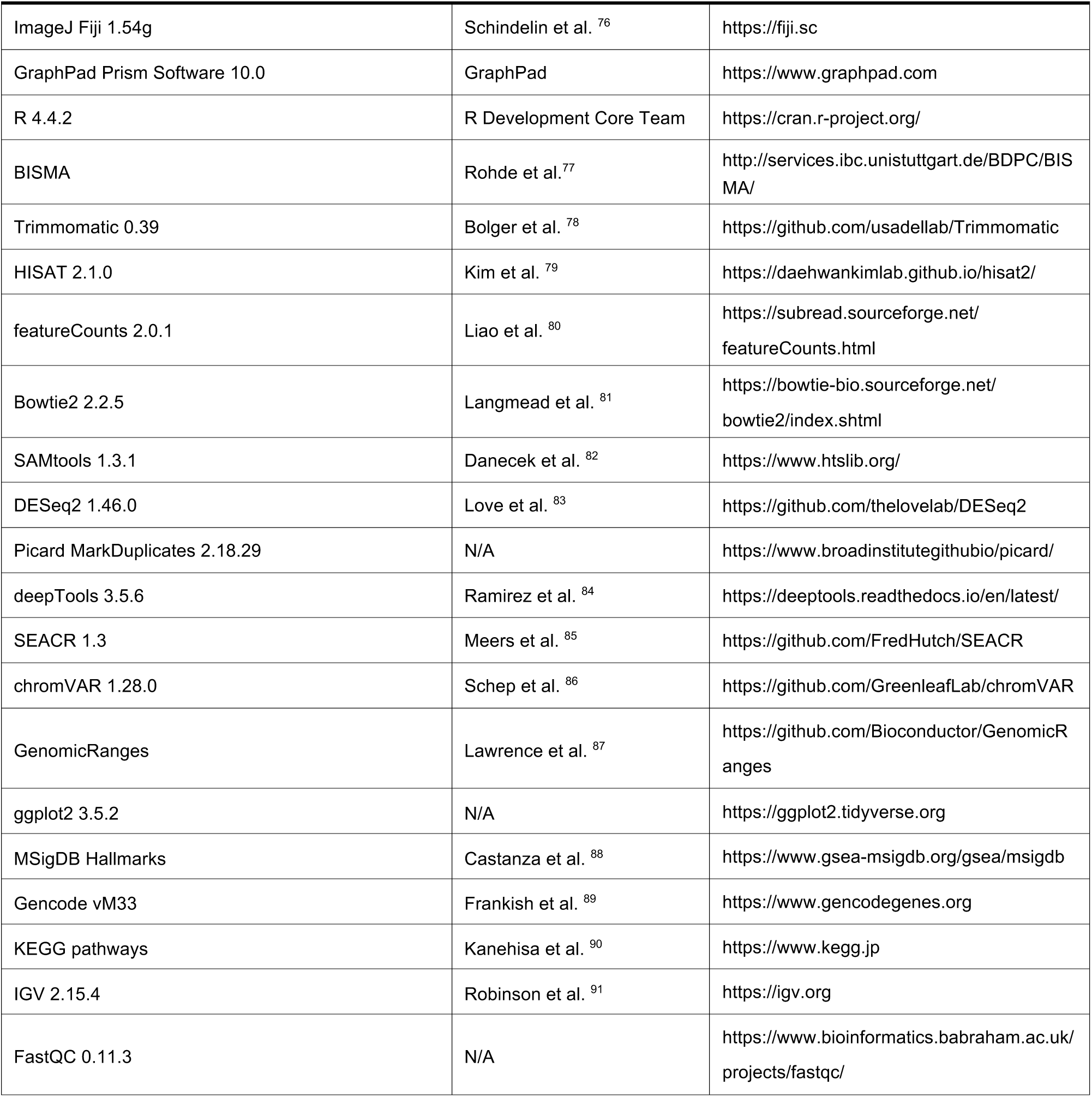

### EXPERIMENTAL MODEL AND STUDY PARTICIPANT DETAILS

#### Mice

Housing, mating and all experimental protocols for mice used in this study were performed in accordance with the guidelines established by the Institutional Animal Care and Use Committee in Guangzhou National Laboratory. Young (3-month-old) and old (24-month-old) C57BL/6 were obtained from Shanghai Model Organisms, Inc., China. Tet2 flox/flox mice were kindly provided by Dr. Guoliang Xu (Shanghai Institute of Biochemistry and Cell Biology, Chinese Academy of Sciences, China). Pax7 CreERT2 mice were purchased from Jackson Laboratory (JAX, stock #017763, USA).

Tet2 flox/flox mice were crossed with Pax7 CreERT2 mice to generate Pax7 CreERT2:Tet2 flox/flox mice. Conditional knockout was induced by Tamoxifen (Sigma, USA) injection as described previously.^92^ In brief, 10mg/ml tamoxifen (Sigma, USA) suspended in corn oil was injected intraperitoneally (ip) into 1-month-old Pax7 CreERT2: Tet2 flox/flox mice for 5 consecutive days at a dose of 100mg/kg body weight per day. Littermates of the same genotype were injected with corn oil as a vehicle control. After 3 months (4-month-old), the mice were sacrificed and whole blood, serum, tibialis anterior (TA), quadriceps (Qu), extensor digitorum longus (EDL), lung, and liver tissues were harvested. Body weight, hair loss ratio, hunchback ratio and muscle force were measured at the day of sacrificing. Running ability was tested a week before sacrificing.

#### Human

Human soleus was obtained from individuals undergoing ligament reconstruction. Fresh muscle tissues were then used for FACS human MuSCs isolation and protein extraction. Ethical approval No: XHEC-D-2019-043.

### METHOD DETAILS

#### Primary Mouse MuSCs Isolation, Expansion, and Differentiation

Primary MuSCs were isolated as previously described.^93^ Briefly, dissected TA muscles were digested with 10ml muscle digestion buffer (DMEM containing 1% penicillin/streptomycin (Hyclone, SV30010, USA), 0.125mg/ml Dispase II (Roche, 04942078001, Germany), and 10mg/ml Collagenase D (Roche, 11088866001, Germany) for 90min at 37°C. The digestion was stopped by adding 2ml of FBS (Hyclone, USA). The digested cells were filtered through 70μm strainers. Red blood cells were lysed by 7ml RBC lysis buffer (0.802% NH_4_Cl, 0.084% NaHCO_3_, 0.037% EDTA, pH7.2-7.4) for 30sec, then filter through 40μm strainers. After staining with antibody cocktails (AF700-anti-mouse Sca-1 (Thermo Fisher Scientific, 56-5981-82, USA), PerCP/Cy5.5-anti-mouse CD11b (BD Biosciences, 550993, USA), PerCP/Cy5.5-anti-mouse CD31 (BD Biosciences, 562861, USA), PerCP/Cy5.5-anti-mouse CD45 (BD Biosciences, 550994, USA), FITC anti-mouse CD34 (BD Biosciences, 553733, USA), and APC-anti-mIntegrin a7+ (R&D Systems, FAB3518A, USA)), the mononuclear cells were subjected for FACS analysis using Influx cell sorter (BD Biosciences, USA). The population of PI^-^CD45^-^CD11b^-^CD31^-^Sca1^-^CD34^+^ Integrin a7^+^ cells were collected. Primary MuSCs were cultured on 0.5mg/ml Type I collagen (Corning, 354249, USA) coated dishes in proliferation medium (F10 medium containing 15% FBS, 5ng/ml IL-1α (PeproTech, 211-11A, USA), 5ng/ml IL-13 (PeproTech, 210-13, USA), 10ng/ml IFN-γ (PeproTech, 315-05, USA) and 10ng/ml TNF-α (PeproTech, 315-01A, USA), 2.5 ng/ml FGF (R&D Systems, 233-FB-025, USA), and 100ug/ml penicillin/streptomycin (Hyclone, SV30010, USA), and differentiated in differentiation medium (DMEM containing 2% horse serum (Hyclone, SH30074.03, USA) and 100ug/ml penicillin/streptomycin (Hyclone, SV30010, USA)) for 48hr as described previously.^4,93^

#### Primary Human MuSCs Isolation

Fresh human muscles were digested similar to that of mouse. After staining with antibody cocktails (APC/Cyanine7 anti-human CD45 (Biolegend, 368516, USA), BV421 anti-human CD56 (BD Biosciences, 562751, USA), PerCP/Cyanine5.5 anti-human CD31 (Biolegend, 303132, USA), AF488 anti-human CD29 (Biolegend, 303016, USA)), the mononuclear cells were subjected for FACS analysis using Influx cell sorter (BD Biosciences, USA). The population of CD31^-^CD45^-^CD29^+^CD56^+^ cells were collected.

#### Running Test

Running studies were conducted as described previously.^65^ In brief, mice were acclimated (run for 9min at 10meters (m)/min followed by 1min at 20 m/min) to the treadmill for 2 consecutive days prior to the experimental protocol. Fed mice were then run for 10 min at 10 m/min followed by a constant speed of 20 m/min until exhaustion.

#### *In Situ* Muscle Force Test

*In situ* hindlimb muscle force analysis was performed with 1300 A 3-in-1 whole animal system (Aurora Scientific, Canada) as previously described.^94,95^ Mice were anesthetized by 3-bromo-2-fluoro-phenyl methanol (31.2 g/kg body weight) ip injection. Mouse TA muscle was exposed by a small incision and half way skin retraction. The distal tendon was carefully severed and sutured to the lever arm of the system, while the ankle was immobilized using clamps. The functional assessment began with three consecutive electrically evoked muscle twitches, followed by a 120sec rest. Subsequently, a series of stimulations at increasing frequencies (10, 20, 30, 40, 50, 60, 80, 100 Hz) were applied to induce tetanic contractions. A 180sec rest was set between each frequency. To assess muscle fatigue, the muscles were subjected to a 500ms, 30 Hz tetanic stimulation every 2sec over a 4.5min period, during which muscle force was monitored. Electrical stimulations were conducted on the common peroneal nerve in the same leg using subcutaneous platinum electrons with 200μs pulse width. The results were analyzed by DMA software (Aurora scientific, Canada).

#### Myofiber Isolation

Myofibers were isolated as described previously.^96^ Briefly, EDL muscles were isolated from tendon to tendon. The isolated EDL was digested in DMEM medium (Gibco, 11965118, USA) contain 0.2% collagenase D (Roche, 11088866001, Germany) for 90min to disassociate myofibers. The digested muscle tissues were triturated three times with a wide bore 5ml pipet. Sit at room temperature for 5min to allow fibers to settle down. Remove the supernatant and wash the myofibers with pre-warmed DMEM medium twice. Pick out single fibers with wide bore yellow tips and applied to RNA extraction, succinate level evaluation, CII activity examination or western blot.

#### *In vivo* Citraconate Treatment

One-hundred microliters of 100mg/mL citraconic acid sodium salt (50 mg/kg body weight; Macklin, C835468, China) or vehicle (PBS) was administered by ip injection to each mouse. Treatment was performed every 1 week for 2 months. TA muscles, lung, and liver samples were collected at day 60 and paraffin-embedded for further analysis.

#### *In vitro* Cell Treatment

MuSCs from 3-month-old oil or tamoxifen injected Pax7 CreERT2: Tet2 flox/flox mice were cultured for one passage, and cells were plated at a density of 1.5 x10^4^ cells/cm^2^ in culture medium. For methyl-succinate treatment, cells were treated with 0.2 mM or 2 mM mono-methyl-succinate (sigma, M81101, USA) for 24hr, then replated in differentiation medium containing the same concentration of mono-methyl-succinate at a density of 4 x10^4^ cells/cm^2^. After 36hr, myotubes were subjected for MitoTracker Red staining, nuclei isolation for immunoblot or RNA isolation for RT-qPCR. For citraconate treatment, cells were treated with 1mM citraconic acid sodium salt (Macklin, C835468, China) for 18hr, then collected for RNA extraction, succinate level evaluation, CII activity examination or metabolic profiling.

#### Histological Analysis

TA muscle, lung, and liver samples were harvested and fixed in 4% paraformaldehyde overnight at 4 ℃. They were then embedded in paraffin and generated 5μm paraffin sections. Tissue sections were stained with hematoxylin and eosin (Sigma-Aldrich, USA) and Masson (Servicebio, G1006, China). For CD45 immunohistological staining, paraffin sections were deparaffinized in xylene, rehydrated with a series of ethanol gradient, incubated in PBS containing 3% H_2_O_2_ for 10min at room temperature to block internal peroxidase activity, then incubated in citrate buffer (0.01M citric acid, pH6), 95 ℃ for 20min to retrieve antigen and blocked in 3% goat serum (Gibco, 16210072, USA) for 60min at room temperature. The sections were incubated in anti-CD45 antibody (1:1000) overnight followed by 3 washes with PBS containing 0.05% Tween 20 at room temperature. They were then incubated in HRP-conjugated secondary antibody followed by 3 washes with PBS containing 0.05% Tween 20 at room temperature, and detected with 3,3′-Diaminobenzidine (DAB) (YEASON, #36303ES01, China). The mounted slides were visualized by EVIDENT VS 200 (Olympus, Japan).

#### Transmission Electron Microscopy

TA muscle was isolated and immediately fixed in 2.5% glutaraldehyde at 4℃ overnight. Washed once with 0.1M phosphate buffer, then fixed in 1% osmium tetroxide solution at room temperature for 1hr, followed by 2 washes using phosphate buffer. The samples were dehydrated using a series of ethanol gradient (30%, 50%, 70%, 80%, 95%, and 100%) and rinsed twice with acetone. The samples were subsequently embedded in 1:1 mix of acetone: Epon812 resin at room temperature for 2hr, followed by embedding in pure Epon812 resin for 24hr. Thin sections (70nm) were sliced using a Leica EM UC7 ultramicrotome and stained with a solution containing 2% uranyl acetate and lead citrate. Images were acquired using a Gatan US1000 2K CCD camera on FEI Tecnai G2 120V electron microscope.

#### Mitochondria Imaging and Function Analysis

For MitoTracker Red staining, live cells were incubated with proliferation or differentiation medium containing 1μM MitoTracker Red (Beyotime, C1035, China) for 30min followed by 3 washes with PBS, then changed to fresh proliferation or differentiation medium. Cells were imaged using the Olympus IXplore SpinSR microscope. Z-stacks were acquired at a 0.5µm step size covering a total depth of 5µm. And images from 10–20 different fields were captured per imaging dish. Raw z-stack images were deconvolved using the Constrained Iterative plug-in and Advanced Maximum Likelihood algorithm was used for analysis of 5 repeats. Maximum intensity projections were generated from deconvoluted z-stack images. Image analysis was conducted using ImageJ Fiji (1.54g, National Institutes of Health, U. S. A.).

#### Seahorse Assay

The Seahorse assay was used to measure OCR (pmol/min) of MuSCs, and isolated mitochondria, as detailed previously described.^97,98^ For MuSCs, 1.5 × 10^4^ cells/well were plated into collagen type I coated Seahorse XF24 plates and cultured overnight to reach more than 90% confluence. The assay medium was prepared by Seahorse XF RPMI Medium contained 10mM Glucose, 1mM Pyruvate, and 2mM L-glutamine, pH7.4. Mitochondrial respiration was monitored at basal state and after sequential injection of the 3µM oligomycin, 6µM FCCP, 2.5µM antimycin A, and 2µM rotenone. For Seahorse assay of mitochondria, the mitochondria from mice MuSCs or hindlimb muscle was isolated using Mitochondria Isolation Kit for Cell (Beyotime, #C3601, China) or Tissue (Beyotime, #C3606, China), respectively. Then 4-6μg mitochondria were placed into each well of Seahorse XF24 plates and centrifuged at 2000g at 4°C for 15 minutes to concentrate the mitochondria to the bottom. To measure mitochondrial OCR in coupling assay, the mitochondria were initially incubated in mitochondrial assay solution (MAS) containing 70mM sucrose, 200mM mannitol, 10mM KH_2_PO_4_, 5mM MgCl_2_, 2 mM HEPES, 1 mM EGTA, and 0.2% (w/v) fatty acid-free BSA, pH7.2, together with 10mM succinate and 2μM rotenone. Mitochondrial respiration was stimulated with the sequential injection of 2mM ADP, 2μM oligomycin, 4μM FCCP, and 4μM antimycin A. In the mitochondria electron transport chain assay, mitochondria were incubated in MAS buffer with 10mM pyruvate, 2mM malate, and 4μM FCCP, and mitochondrial respiration was stimulated with the sequential injection of 2μM rotenone, 10mM succinate, 4μM antimycin A, 10mM ascorbate, and 0.1mM TMPD.

#### Metabolite Analyses by Mass Spectrometry

Mass spectrometry was used to measure metabolites of MuSCs and tissue mitochondria. For MuSCs, organic acids were analyzed at LipidALL Technologies. In brief, metabolites were extracted from MuSCs using acetonitrile: water (1:1) and derivatized with 3-Nitrophenylhdyrazine.^99^ Metabolites were analyzed on a Jasper HPLC coupled to Sciex 4500 MD system. In brief, individual metabolites were separated on a Phenomenex Kinetex C18 column (100 x 2.1mm, 2.6µm) using 0.1% formic acid in water as mobile phase A and 0.1% formic acid in acetonitrile as mobile phase B. d4-succinic acid, d4-citric acid, d3-malic acid, 13C3-latic acid, d3-pyruvic acid, d4-fumaric acid purchased from Cambridge Isotope Laboratories were used as internal standards for quantitation. For metabolites of tissue mitochondria, hindlimb muscles from 4-month-old male WT and Tet2 KO mice were collected. Mitochondria were then isolated using Mitochondria Isolation Kit for Tissue and nitrogen frozen. Then samples were prepared and analyzed on LC/MS platforms based on a Thermo Fisher Vanquish UHPLC equipped with an ACQUITY UPLC BEH Amide column (1.7 μm, 2.1 mm x 100 mm, Wasters) and a Thermo-TSQ Vantage™ mass spectrometer. in Shanghai Bioprofile company (China). Raw MRM data files were processed by peak finding, alignment, and filtering using Xcalibur Qual browser software (Thermo Fisher Scientific, USA). Simcap 14 software (Umetrics, Umeå, Sweden) was used for all multivariate data analysis and modeling.

#### Metabolite and Enzyme Activities Analysis

Succinate and the activity of SDH/Complex II were measured using Succinate Assay Kit (Colorimetric, USA) (Abcam, ab204718, USA) and Complex II Enzyme Activity Microplate Assay Kit (Colorimetric, USA) (Abcam, ab109908, USA) based on DCPIP method, respectively. All colorimetric assays were performed according to the manufacture’s protocol.

#### Enzyme-linked Immunosorbent Assays (ELISA)

The levels of CCL6 and CXCL11 in mouse serum were measured using Mouse CCL6 SimpleStep ELISA Kit (Abcam, ab277715, USA) and Mouse CXCL11 SimpleStep ELISA Kit (Abcam, ab204519, USA), respectively. All ELISA assays were performed according to the manufacture’s instruction.

#### Immunoblotting

Proteins were separated by 10% SDS-PAGE according to standard protocols, transferred to Immobilon-P PVDF Membrane (Millipore, IPVH00005, USA). After blocking with 5% skim milk in TBST, membranes were incubated with the primary antibodies: H4K31succ (1:3000), H4 (1:1000), Flag (1:4000), beta-Actin (1:3000), overnight at 4 ℃. After washing in TBST, horseradish peroxidase-conjugated anti-rabbit or anti-mouse IgG antibody were applied. The membranes were washed 3 times with TBST and imaged by MiniChemi 580 (Beijing Sage Creation Science Co., Ltd, China).

#### RT-qPCR

Total RNA was extracted by TRIzol Reagent (Thermo Fisher Scientific, 15596-018, USA), and reverse transcription was performed using Reverse-transcriptase M-MuLV (NEB, M0253L, USA), followed by qPCR analysis using SYRB Green qPCR Mix by QuantStudio6 Flex (Thermo Fisher Scientific, USA). The expression level of each gene was normalized to that of *Gapdh* using delta-delta Ct method. Primers used were as follows: m*Tet2* (forward, 5’-CAATAATCAGGTTGAATTTGAAC-3’, reverse, 5’-GAGGGTGA CCACCACTGTACTGC-3’), m*Acod1* (forward, 5’-GGTATCATTCGGAGGAGCAAG AG-3’, reverse, 5’-ACAGTGCTGGAGGTGTTGGAAC-3’), m*Hdac11* (forward, 5’-AAGGCATCTCCAGAGCCACCAT-3’, reverse, 5’-CAGGGTAGATGTGGCGGTTG TA-3’), m*Sirt5* (forward, 5’-ATCGCAAGGCTGGCACCAAGAA-3’, reverse, 5’-CTA AAGCTGGGCAGATCGGACT-3’), m*Sirt7* (forward, 5’-CTGGAGATTCCTGTCTACAACCG-3’, reverse, 5’-AGTGACTTCCTACTGTGGCTGC -3’), m*Cpt1a* (forward, 5’-GGCATAAACGCAGAGCATTCCTG-3’, reverse, 5’-CAGTGTCCATCCTCTGAGT AGC-3’), m*Kat2a* (forward, 5’-CACGGAAATCGTCTTCTGTGCC-3’, reverse, 5’-CG TACTCGTCAGCATAGGTGAG-3’), m*Ccl2* (forward, 5’-TGGAGCATCCACGTGT TGGC-3’, reverse, 5’-TGGGACACCTGCTGCTGGTG-3’), m*Ccl5* (forward, 5’-CCTGCTGCTTTGCCTACCTCTC-3’, reverse, 5’-ACACACTTGGCGGTTCCTTCGA - 3’), m*Ccl6* (forward, 5’-CCAGGCTGGCCTCATACAAG-3’, reverse, 5’-TTACATGG GATCTGTGTGGC-3’), m*Ccl8* (forward, 5’-CCTGCTGCTCATAGCTGTCC-3’, reverse, 5’-CAGCTTCCATGGGGCACTGG-3’), m*Ccl11* (forward, 5’-TTCCTGCTGCTCA CGGTCAC-3’, reverse, 5’-TGTAGCTCTTCAGTAGTGTG-3’), m*Ccl12* (forward, 5’-TGCTCATAGCTACCACCATC-3’, reverse, 5’-TTGTGATTCTCCTGTAGCTC-3’), m*Ccl24* (forward, 5’-CTCTGCTACGATCGTTGCCG-3’, reverse, 5’- TGGCCAACTG GTAGCTAACC-3’), m*Cxcl5* (forward, 5’-TTCCATCTCGCCATTCATGC -3’, reverse, 5’-CAACGCAGCTCCGTTGCGGC-3’), m*Cxcl10* (forward, 5’-CTGCCTCATCCTGC TGGGTC-3’, reverse, 5’-AGGATAGGCTCGCAGGGATG-3’), m*Cxcl11* (forward, 5’-GCCCTGGCTGCAATATCTGG-3’, reverse, 5’-TTTGTCGCAGCCGTTACTCG-3’), m*Cxcl12* (forward, 5’-CAAGGTCGTCGCCGTGCTGG-3’, reverse, 5’-GATGCTTGAC GTTGGCTCTG-3’), m*IL6* (forward, 5’-CCACTTCACAAGTCGGAGGC-3’, reverse, 5’-TGTATCTCTGGAAGTTTCAG-3’), m*IL15* (forward, 5’-GTAGGTCTCCCTAAAACAGAGGC-3’, reverse, 5’-TCCAGGAGAAAGCAGTTCATTGC-3’), m*IL1b* (forward, 5’-TGGACCTTCCAGGATGAGGACA-3’, reverse, 5’-GTTCATCTCGGAGCCTGT AGTG-3’), m*Tnfsf11* (forward, 5’-GTGAAGACACACTACCTGACTCC-3’, reverse, 5’-GCCACATCCAACCATGAGCCTT-3’), m*Tnf* (forward, 5’-GGTGCCTATGTCTCAGCCTCTT-3’, reverse, 5’-GCCATAGAACTGATGAGAGGGAG-3’), m*Gapdh* (forward, 5’-GCCAGCCTCGTCCCGTAGACA-3’, reverse, 5’-CAACAATCTCCACTTTGCCAC TGC-3’), h*TET2* (forward, 5’-GCTTACCGAGACGCTGAGGAAA-3’, reverse, 5’-AG AGAAGGAGGCACCACAGGTT-3’), h*GAPDH* (forward, 5’-GTCTCCTCTGACTTCAACAGCG-3’, reverse, 5’-ACCACCCTGTTGCTGTAGCCAA-3’).

#### Bisulfite Sequencing

Genomic DNA was extracted and treated with EpiArt DNA Methylation Bisulfite Kit (Vazyme, EM101, China) according to the manufacturer’s instructions. Bisulfite-treated DNA was subjected to nested PCR amplification, then purified with the Gel Extraction Kit (TianGen, DP204, China) and cloned into T-Vector pMD20 (Takara, 3270, Japan). Individual clones were sequenced by standard Sanger sequencing. Data were analyzed by BISMA (http://services.ibc.uni-stuttgart.de/BDPC/BISMA/).^77^ Primers used were as follows: m*Hdac11*-DMR1-1 (forward, 5’-TTATTGAGATATAATTTTAGTTTTTAGATTGT-3’, reverse, 5’-ATCCCACAATAACCTAAATTTATCTCCAACTC-3’), m*Hdac11*-DMR1-2 (forward, 5’-GAGTTGGAGATAAATTTAGG TTATTGTGGGAT-3’, reverse, 5’-CTAACTTTACCACAACCAAAACTACCCATCCC-3’), m*Hdac11*-DMR2-1 (forward, 5’-GGGGAGGGGGAGTGATGGGGGGGGTGTTGTTG-3’, reverse, 5’-ACACAACACCCACAAAAAAACCCAAAAAACACC-3’), m*Hdac11*-DMR1-1 (forward, 5’-GGGAGGAAAGGGTTTATTTAGTTTATATTTTT-3’, reverse, 5’-TATC AACTTAACACAAACTAAAATTATCACAA-3’).

#### ChIP-qPCR

ChIP assays were performed as previously described.^100,101^ Briefly, crosslinking was performed with 1% formaldehyde for 10min at room temperature, then stopped by 125mM glycine for 5min. Nuclei were isolated in cell lysis buffer (50mM HEPEs, pH7.6, 10mM KCl, 1.5mM MgCl_2_, 1mM EDTA, 0.5mM EGTA, 0.5% Triton X-100, 1mM PMSF, 1mM DTT) and chromatin was further extracted from the nuclei using nuclei lysis buffer (50mM HEPEs pH7.6, 1mM EDTA, 0.5mM EGTA, 1% Triton X-100, 0.1% deoxycholate, 1mM PMSF, 1mM DTT). The chromatin was sheared to 200∼500bp by sonication using Bioruptor PICO (Diagenode SA, Belgium) and incubated with antibodies. Dynabeads^TM^ Protein G (Invitrogen, 10004D, USA) were used to capture the precipitated chromatin by antibody. The beads were washed sequentially with sonication buffer (20mM HEPEs, pH8, 150 mM NaCl, 2mM EDTA, pH8, 0.25% SDS and 1% Triton X-100), high-salt buffer (20mM HEPEs, pH8, 500mM NaCl, 2mM EDTA, pH8, 0.1% SDS and 1% Triton X-100), LiCl buffer (10mM HEPEs, pH8, 250mM LiCl, 1mM EDTA, pH8, 1% NP-40), TE buffer (1mM EDTA, pH8, 10mM HEPEs, pH8) and the crosslink was reversed by incubating in 200mM NaCl, 125μM proteinase K, and 62.5μg/ml RNase A at 65 ℃ overnight. The DNA fragments were purified by phenol-chloroform extraction and detected by qPCR. The fold enrichment was calculated against IgG ChIP-qPCR. Primers used were were listed below. m*Acod1*-TSS (forward, 5’-AAGTGAGGTCATTGGGGAGG-3’, reverse, 5’-CGCCAT CAAGTGCTACGATAC-3’), Background (forward, 5’-CATTCCCAGAAGCTGTT TCATG-3’, reverse, 5’-GGGATGGCCTGTCCCAGCTAGG-3’), m*Hdac11*-DMR1-1 (forward, 5’-CGTGCAGATAGACAGGCTTCTG-3’, reverse, 5’-CTGAGTTTATC TCCAGCTCCCG-3’), m*Hdac11*-DMR1-2 (forward, 5’-TCCTCTGGATCTCTCAG TCTGC-3’, reverse, 5’-TGCTCAGCAGTGGCTCCTCTAC-3’), m*Hdac11*-DMR2-1 (forward, 5’-CGCTCAGGTGAGACCTATCTAA-3’, reverse, 5’-GACCCAAGAAAC ACCACCGTGG-3’), m*Hdac11*-DMR2-2 (forward, 5’-AGGTCTCTTTAGGGTTCCCA CG-3’, reverse, 5’-AACGATGTGGAAGTGTAAGCTG-3’), m*Ccl5* (forward, 5’-GTCTGCCTGAGCTGCAGAGC-3’, reverse, 5’-CTGCAGTGAGGATGATGGTG-3’), m*Ccl6* (forward, 5’-CAGGCCAGAGAATCAGGGAG-3’, reverse, 5’-CCTTGTATGA GGCCAGCCTG-3’), m*Ccl12* (forward, 5’-GTCAGATGGAACTTCCAGGG-3’, reverse, 5’-TGGTCCAGCCAATACCTGAG-3’), m*Cxcl10* (forward, 5’-GCACTCGAGAAGC GCTTCAT-3’, reverse, 5’-GGTGTCCCTTACCTTGAGTC-3’), m*Cxcl11* (forward, 5’-CCAGGCAGAGAGCTGCAGCG-3’, reverse, 5’-GGAACAGACAGACTCAGAAG-3’), Gene desert (forward, 5’-TCCTCCCCATCTGTGTCATC-3’, reverse, 5’-GGATCCAT CACCATCAATAACC-3’).

#### CUT&Tag sequencing Library Construction

CUT&Tag-seq of FACS sorted MuSCs was performed using Hyperactive® Universal CUT&Tag Assay Kit for Illumina (Vazyme, TD903, China). Briefly, 5 x 10^4^ cells were incubated with ConA beads for 10min at room temperature. Primary antibodies, H4K31succ (1:50), were then added and incubated with the cells-beads complex overnight at 4℃. Secondary antibodies were added and incubated for 45min at room temperature. After washing with Dig-wash Buffer, pA/G-Tnp was added into each tube and incubated for 1hr at room temperature. Tagmentaion buffer containing Spike-inAmpR prepared as previously described^102^ was then added to each CUT&Tag reaction and incubated for 1hr at 37 ℃. At last, DNA was eluted and library was prepared using TruePrep® Index Kit V2 for Illumina (Vazyme, TD202, China) according to the manufacturer’s instructions.

#### CUT&Tag-seq Data Pre-processing

These libraries were sequenced by Illumina NovaSeq 6000 instrument (Illumina, USA) with 150bp reads and paired-end parameter by Annoroad Gene Technology company, China. The 150 bp paired-end CUT&Tag-seq reads were trimmed using Trimmomatic_0.39^78^ and aligned to the mouse reference genome (GRCm39) or Spike-in sequence^102^ using Bowtie2^81^ with parameters -X 700 -no-mixed -no-discordant -no-unal - local -very-sensitive-local. Aligned reads are filtered for uniquely aligned reads with the Samtools^82^ flag -q 10 and duplicate reads was removed by using the Picard Mark-Duplicates (Picard, USA) (https://www.broadinstitutegithubio/picard/). Per-sample normalization factors were calculated as previously described^102^ using reads mapped to the Spike-in sequence, and used to generate normalized bigwig and bedgraph files using bamCoverage from deepTools.^84^ Peak regions of each sample were identified by SEACR_1.3^85^ with options -stringent. For each peak, fragment counts were obtained using R chromVAR^86^ and normalized using normalization factors, then log2 fold enrichment was calculated as log2(counts (H4K31succ) / counts (IgG)). Peaks with log2 fold enrichment greater than 2.5 in either sample were used for downstream analysis.

#### Cut&tag, ChIP, and ATAC sequencing Data Analysis

The CUT&Tag, ChIP^69-71^ or ATAC^4^ intensity for the gene bodies (transcriptional start site to transcriptional termination sties as annotated in GRCm39) of all genes or peaks region was calculated using the computeMatrix from deepTools.^84^ Composite metagene plots were generated by plotProfile from deepTools.^84^ Heatmaps were generated using plotHeatmap from deepTools^84^ with parameters–sortUsing sum, such that sites with the largest signal are oriented at the top of the heatmap, and sites with the smallest signal are oriented at the bottom of the heatmap.

For each H4K31succ peak region in WT MuSCs, fragment counts for H3K4me3, H3K27ac, H3K4me1, H3K36me3, H3K27me3 were obtained from published ChIP-seq datasets^69-71^ and log2 fold enrichment (relative to Input) was calculated. H4K31succ peaks located within promoter regions (log2 (H3K4me3/Input)>0.4 and log2 (H3K27ac/Input)>0.4) were designated as type 1. H4K31succ peaks located within enhancer regions (log2 (H3K4me1/Input)>0.2 and log2 (H3K27ac/Input)>0.2) were designated as type 2. H4K31succ peaks located within gene bodies (log2 (H3K36me3/Input)>0.2) were designated as type 3. H4K31succ peaks having only H3K27me3 marks (log2 (H3K27me3/Input)>0.2) were designated as type 4. H4K31succ peaks having none of these histone marks were designated as type 5. These five H4K31succ peak types were used for downstream analysis. Heatmaps of relative CUT&Tag-seq and ChIP-seq signal by type were generated based on method described above. Sites were ordered based on type assignment as indicated. Sites within each type were ranked in descending order.

Peak annotation was performed by intersecting peak.bed file with gff file (Gencode vM33) using bedtools.^103^ Biotypes were derived from Gencode vM33 annotations for mouse assembly GRCm39. H4K31succ peaks were annotated relative to genomic features as follows: peaks with their center located within 0-2 kb upstream of a gene were designated as TSS_0-2kb, peaks with their center located within gene bodies (transcriptional start site to transcriptional termination sties) were designated as Gene body, peaks with their center located within intergenic regions were designated as Intergenic.

H4K31succ peaks from two conditions (WT vs Tet2 KO) were first merged to generate a master peak list using R GenomicRanges.^87^ For each peak in the list, log2 fold enrichment was calculated separately for each condition. Peaks with log2 fold enrichment difference between Tet2 KO and WT >1.2 were designated as Gained. Peaks with log2 fold enrichment difference between -1.2 and 1.2 were designated as Maintained. Peaks with log2 fold enrichment difference < -1.2 were designated as Lost. These three differential peak types were used for downstream analysis. For accessibility changes analysis upon Tet2 KO of these three peak types, ATAC signal of WT or Tet2 KO MuSCs were obtained from our previously published dataset.^4^ For expression changes analysis upon Tet2 KO of genes overlapping with these three peak types, RNA-seq data were obtained from our previously published dataset.^4^ Genes overlapping with Gained or Lost peaks were used for further functional enrichment analysis. Differential analysis of H4K31succ CUT&Tag-seq in young or old MuSCs were performed in the same way.

Genes with gained or lost (defined as described above) H4K31succ peaks upon Tet2 KO or ageing were used for functional enrichment analysis, which was performed using our self-defined functional terms (Table S1) generated by combining MSigDB hallmarks and KEGG pathways. Bubble plots were generated using R ggplot2, P-values were calculated using the phyper function in R (4.4.2).

For visualization of selected genome regions, we used IGV-2.15.4^104^ with the bigwig files generated by bamCoverage^84^ using genome build GRCm39.

#### RNA-seq Library Preparation and Data Processing

RNA-seq of TA muscle, lung, liver from oil or tam injected Pax7 CreERT2: Tet2 flox/flox mice was performed as follows: total RNA was extracted by TRIzol Reagent (Thermo, 15596-018, USA), 1μg of total RNA were used for preparing RNA-seq libraries using TruSeq RNA Library Preparation Kit v2 (Illumina, RS-122-2001, USA). Libraries were sequenced on an Illumina HiSeq 2000 (Illumina, USA) to generate 150 base paired-end reads. The raw RNA-seq reads were first evaluated by FastQC (https://www.bioinformatics.babraham.ac.uk/projects/fastqc/) and preprocessed by Trimmomatic_0.39^78^ to trim sequencing reads, eliminating Illumina adaptor remains, and to discard reads that were shorter than 30 bp. The resulting reads were mapped against reference genome GRCm39 using Hisat2 v.2.1.0^79^ with default parameters. FeatureCounts v.2.0.1^80^ was used to generate a matrix of mapped fragments per RefSeq annotated gene, with annotations from Gencode vM33. Differentially expressed genes were found using R package DESeq2.^83^ Genes with adjusted p-value < 0.05 were considered for further analysis. Functional enrichment was performed using our self-defined functional terms (Table S1). Bubble plots were generated using R ggplot2, p-values were calculated using the phyper function in R (4.4.2). For comparative enrichment analysis of Tet2 KO cells/tissues and previously published aging datasets: qMuSCs, 1-DPI MuSCs, 3-DPI MuSCs, skeletal muscle, lung, and liver,^54,72-74^ differential expression analysis of all published dataset was first performed using R package DESeq2, then functional enrichment was performed as described above.

#### Analysis of Publicly Available Data

RNA-seq data, WBGS data, and ATAC-seq data of WT or Tet2 KO MuSCs were from our previously published work and processed as described in the paper.^4^ ChIP-seq data of H3K4me3, H3K36me3, or H3K27me3 from MuSCs,^69^ H3K4me1 from C2C12,^70^ and H3K27ac from C2C12^71^ were downloaded from NCBI Gene Expression Omnibus GSE47362, GSE25308 and GSE37525, respectively, and aligned to GRCm39 according to the parameters described.^4^ Replicates were merged and converted to bigwig format for heatmap analysis. RNA-seq data of myofibers stripped of MuSCs from 4-month-old (young) or 22-month-old (old) mice^62^ were downloaded (GSE193261) as FASTQ files and mapped according to the parameters described above. RNA-seq data of HFSCs isolated from young or old mice^55^ were downloaded (GSE148619) as FASTQ files and mapped according to the parameters described above. RNA-seq data of qMuSCs, FiMuSCs, aMuSCs isolated from young or old mice^59^ were downloaded (GSE113631) as FASTQ files and mapped according to the parameters described above. Raw counts for RNA-seq data of TA muscle from 6-month-old (young), 21-month-old (old), 24-month-old (old), 27-month-old (old) mice^72^ were downloaded from GSE226117. Raw counts for RNA-seq data of liver from 3-month-old (young) and 29-month-old (old) mice^74^ were downloaded from NCBI BioProject database under accession number PRJNA281127. Differential gene expression and functional enrichment analysis was performed as described above. Processed data for RNA-seq of qMuSCs, 1-DPI MuSCs, 3-DPI MuSCs isolated from young or old mice^54^ were downloaded from GSE183643. Functional enrichment analysis was performed as described above. Processed data for lung RNA-seq from young or old mice^73^ were downloaded from GSE124872. Processed data for HSCs isolated from young or old mice were downloaded from GSE47819. Processed data for NSCs isolated from young or old mice were downloaded from GSE101610^57^ and a previously published paper.^58^ Processed data for skeletal muscle RNA-seq from young or aged people^75^ were downloaded from GSE167186.

#### Statistical Analysis

Box plots, dot plots, or points & connecting line with error bars were plotted in GraphPad Prism 10.0 (GraphPad Software, San Diego, CA, USA) and have a line at the median, and whiskers show the standard deviation unless noted otherwise. Statistical differences between groups were determined by unpaired two-tailed t-test in Prism. Statistical significance was set at p < 0.05. ns indicated no significant difference, * indicated p < 0.05, ** indicated p <0.01, *** indicated p < 0.001, **** indicated p < 0.0001. The numbers of biological replicates and technical repeats in each experimental group were indicated in figure legends.

---

**Figure S1.**
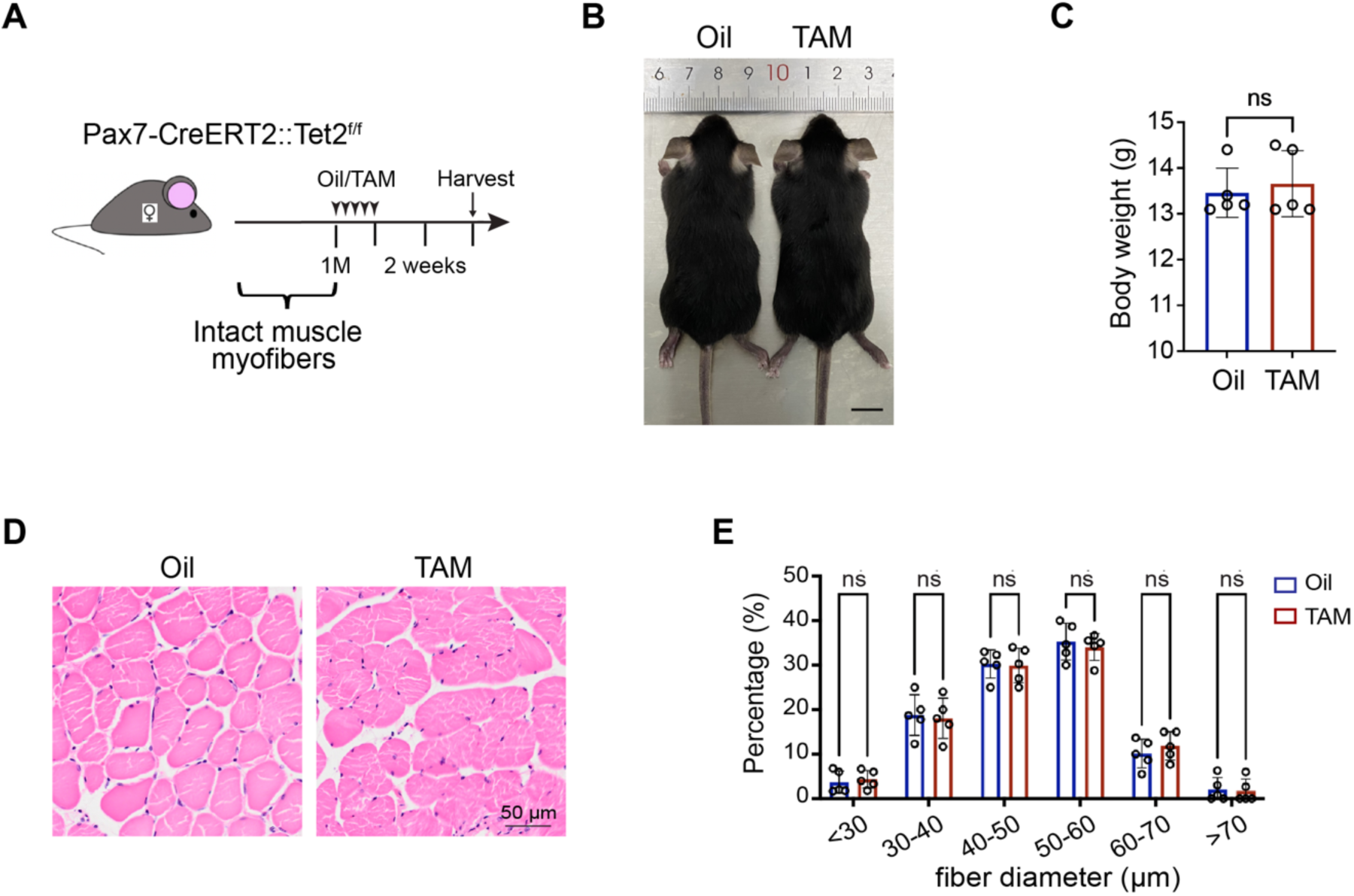
No obvious phenotypes shown in MuSC specific *Tet2* KO mice 2 weeks post ablation induction. Related to Figure 1. **(A)** Scheme of MuSC specific *Tet2* KO induction. **(B)** Representative images of MuSC specific *Tet2* KO mice 2 weeks post KO induction. Scale bar: 1cm. **(C)** Statistical analysis of body weights of *Tet2* KO mice (n = 5 in each group). **(D)** Representative images of H&E staining of TA muscles (n = 5 in each group). Scale bar, 50μm. **(E)** Quantification of TA myofiber sizes. **(C) and (E)** Data were shown in mean ± SD form. Data were analyzed by unpaired t-test. ns indicated not significant.

**Figure S2.**
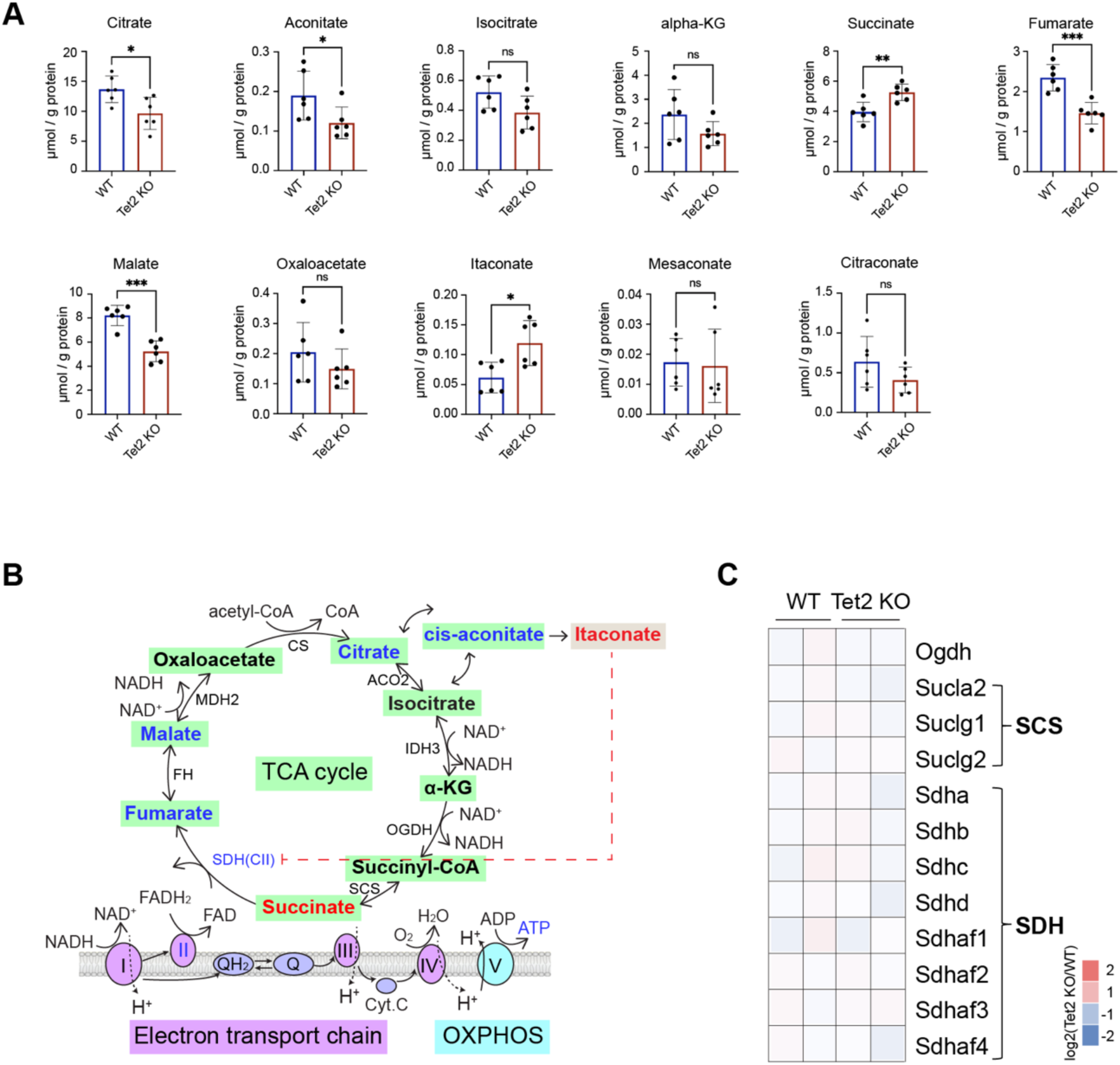
SDH defects in *Tet2* KO MuSCs leads to succinate accumulation. Related to Figure 2. **(A)** Quantification of metabolites in WT and *Tet2* KO MuSCs detected by targeted LC-MS/MS analysis (n=6 in each group). **(B)** Illustration of TCA cycles. Blue color indicated metabolites and enzymes decreased in *Tet2* KO MuSCs. Red color indicated metabolites and enzymes increased in *Tet2* KO MuSCs. Data were analyzed by unpaired t-test based on 3-6 independent experiments. **(C)** Heatmap illustrating the log2 fold change of genes involved in synthesis (*Ogdh*, SCS) and dehydrogenation (SDH) of succinate in *Tet2* KO compared to that in WT MuSCs. RNA-seq data were from our previously published dataset: GSE158649.

**Figure S3.**
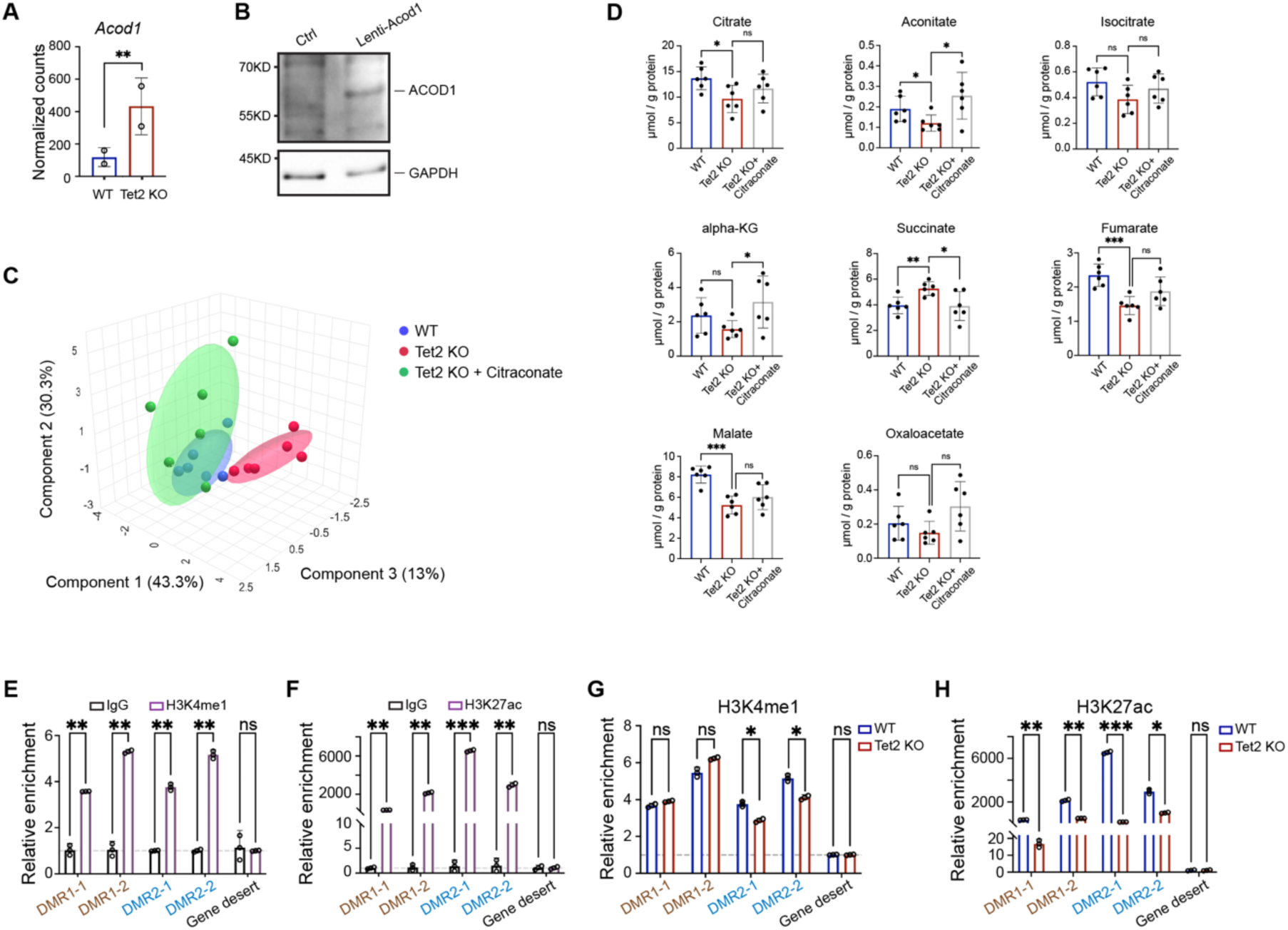
Disruption of a Tet2-Hdac11-Acod1 axis results in succinate accumulation in *Tet2* KO MuSCs. Related to Figure 3. **(A)** Normalized sequencing counts of *Acod1* in WT and *Tet2* KO MuSCs. RNA-seq data were from our previously published dataset: GSE158649. **(B)** Immunoblotting analysis of ACOD1 expression in control (Ctrl) and *Acod1* overexpressing (lenti-Acod1) MuSCs, respectively. GAPDH served as a loading control. **(C)** Scatter plot of targeted LC-MS/MS analysis of WT MuSCs, *Tet2* KO MuSCs, and *Tet2* KO MuSCs treated by citraconate. The plot was generated by MetaboAnalyst 6.0. **(D)** Quantification of targeted LC-MS/MS analysis of metabolites in WT, *Tet2* KO, and *Tet2* KO MuSCs treated by citraconate (n=6 in each group). **(E)-(F)** Quantification of H3K4me1 (E) and H3K27ac (F) enrichment in DMR1 and DMR2 of *Hdac11* by ChIP-PCR (n=3 in each group). **(G)-(H)** Quantification of H3K4me1 (G) and H3K27ac (H) enrichment in DMR1 and DMR2 of *Hdac11* in *Tet2* KO MuSCs (n=3 in each group). **(A) and (D)-(H)** Data were shown in mean ± SD form. Data were analyzed by unpaired t-test. * indicated *p* < 0.05; ** indicated *p* < 0.01; *** indicated *p* < 0.001; ns indicated not significant.

**Figure S4.**
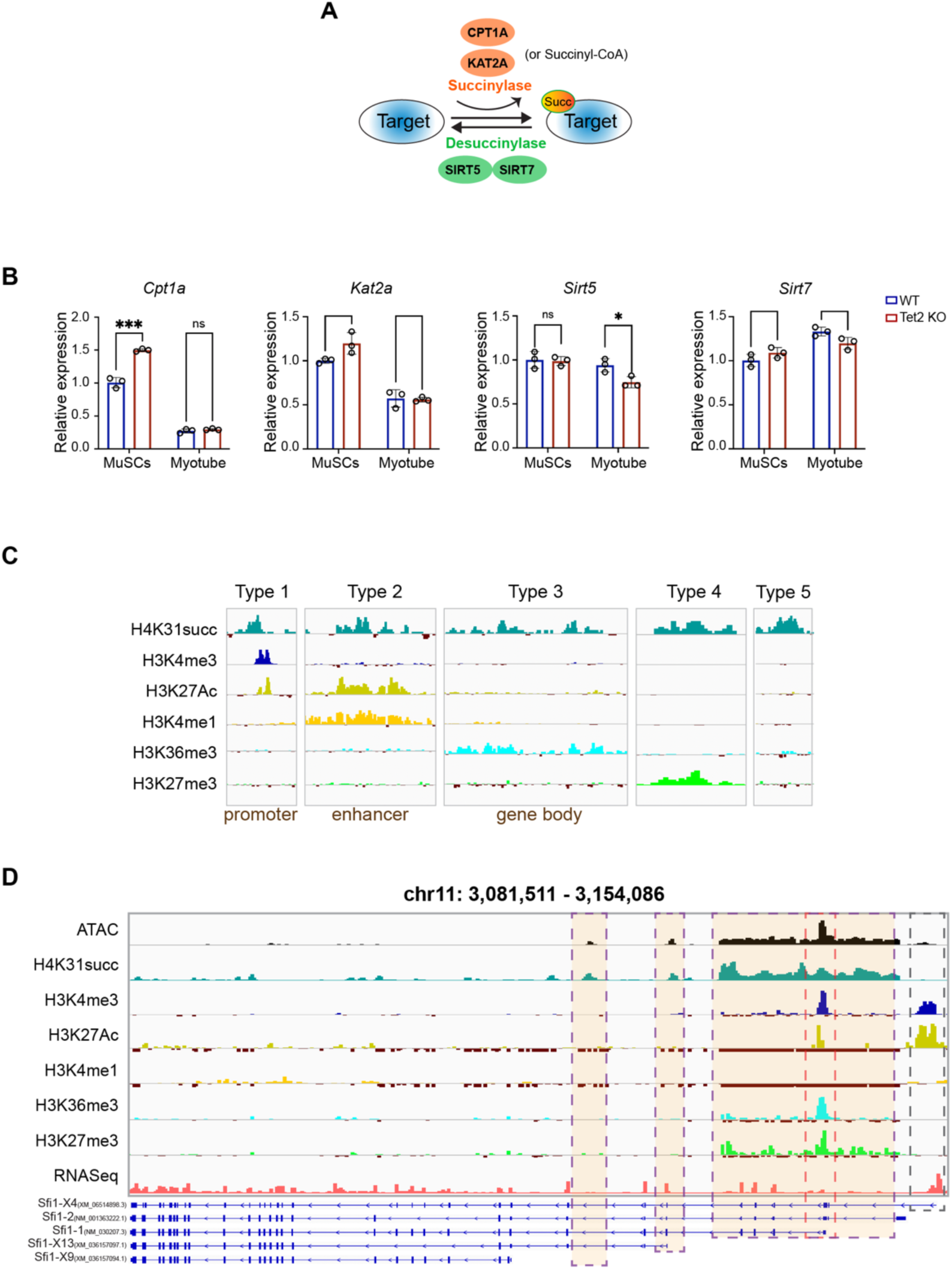
H4K31succ is an active histone mark. Related to Figure 4. **(A)** Illustration of succinylation and desuccinylation reactions. **(B)** Relative expression levels of genes involved in succinylation (*Cpt1a*, *Kat2a*) and desuccinylation (*Sirt5*, *Sirt7*) in WT and *Tet2* KO MuSCs and myotubes, respectively (n=3). Data were shown in mean ± SD form. Data were analyzed by unpaired t-test based on 3 independent experiments. *** indicated *p* < 0.001; ns indicated not significant. **(C)** IGV browser snapshots showing the distribution of reads for the indicated histone modifications around five representative H4K31succ peak types. **(D)** Genome browser images of the representative canonical promoter peak (with high H3K4me3 and H3K27ac signal, grey dashed box), type 5 H4K31succ peaks with high chromatin accessibility (purple dashed boxes) and type 1 H4K31succ peak (pink dashed box) at the promoter of *Sfi1* isoforms in WT MuSCs.

**Figure S5.**
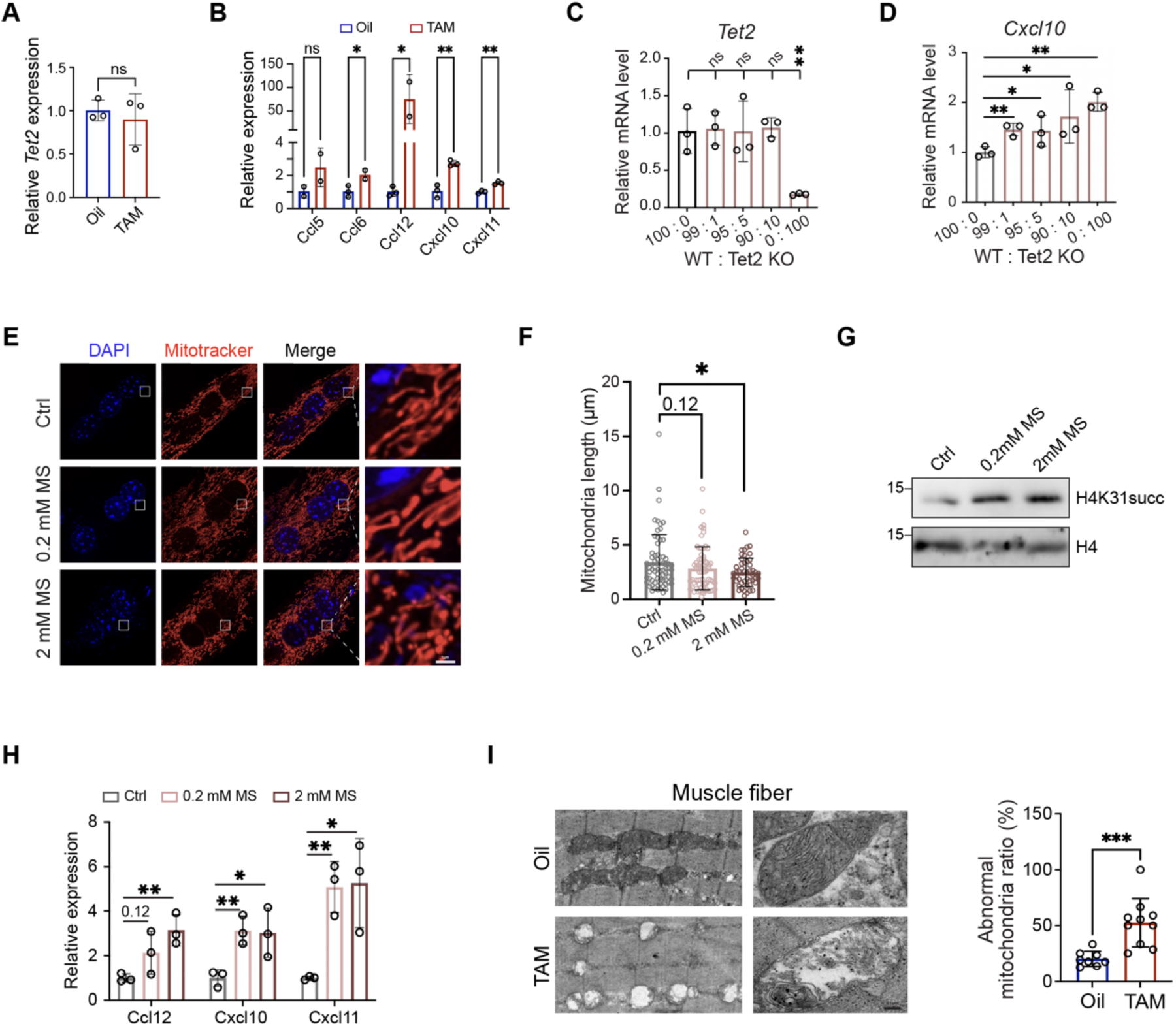
Succinate treatment results in mitochondria impairment and increases of inflammation in myotubes. Related to Figure 5. **(A)-(B)** Relative expression level of *Tet2* (A) *and Ccl5*, *Ccl6*, *Ccl12*, *Cxcl10*, *Cxcl11* (B) in single myofibers isolated from EDL of oil or TAM injected Pax7 CreERT2:Tet2 f/f mice (n=3 in each group). **(C)-(D)** Analysis of relative expression level of *Tet2* (C) and *Cxcl10* (D) in WT myotubes incorporated with various ratio of Tet2 KO MuSCs (n=3 in each group). **(E)** Representative images of the mitochondria stained with MitoTracker (red) and DAPI (blue) in WT myotubes treated with 0mM (Ctrl), 0.2mM, and 2mM mono-methyl-succinate (MS). Scale bar, 1μm. **(F)** Quantification of the mitochondria length in panel E. **(G)** Immunoblotting of H4K31succ in WT myotubes treated with 0 mM (Ctrl), 0.2mM, and 2mM mono-methyl-succinate (MS). H4 served as a loading control. **(H)** Analysis of relative expression level of *Ccl12*, *Cxcl10*, and *Cxcl11* in WT myotubes treated with 0mM (Ctrl), 0.2mM, and 2mM mono-methyl-succinate (MS) by RT-qPCR (n=3 in each group). **(I)** Representative TEM images of mitochondria (left) in skeletal muscle from oil or TAM injected Pax7 CreERT2:Tet2 f/f mice, and quantification of the ratio of abnormal mitochondria (right) (n=10). Scale bar, 0.1µm. **(A)-(D), (F), and (H)-(I)** Data were shown in mean ± SD form. Data were analyzed by unpaired t-test. * indicated *p* < 0.05; ** indicated *p* < 0.01; *** indicated *p* < 0.001; ns indicated not significant.

**Figure S6.**
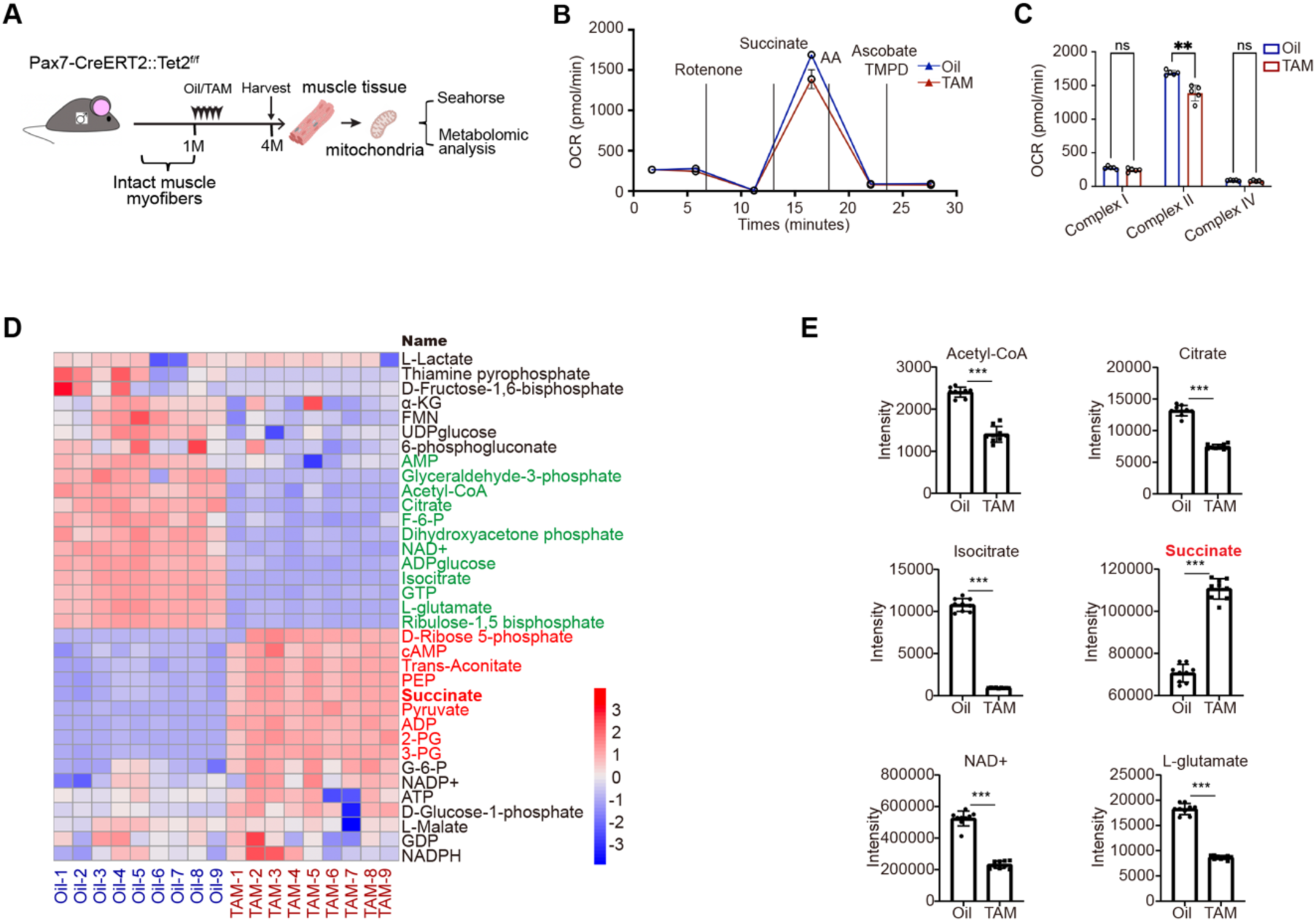
Succinate accumulates in skeletal muscles from *Tet2* KO mice. Related to Figure 6. **(A)** Scheme of experiments. **(B)** Seahorse respiratory complex assays showing normalized OCRs in isolated mitochondria from oil or TAM injected *Pax7* CreERT2: *Tet2* f/f mice. Data were normalized to protein level. **(C)** Quantification of respiration of each mitochondrial ETC complex from oil or TAM injected *Pax7* CreERT2: *Tet2* f/f mice (n=5 in each group). **(D)** Heatmap analysis of LC/MS metabolomics data. The metabolites are color-coded to represent their relative levels in TAM (*Tet2* KO) compared to Oil (Ctrl) injected *Pax7* CreERT2: *Tet2* f/f mice. **(E)** Quantification of the TCA metabolites showing significant changes in TAM injected *Pax7* CreERT2: *Tet2* f/f mice (*Tet2* KO) (n=9 in each group). **(C) and (E)** Data were shown in mean ± SD form. Data were analyzed by unpaired t-test. ** indicated *p* < 0.01; *** indicated *p* < 0.001; ns indicated not significant.

**Figure S7.**
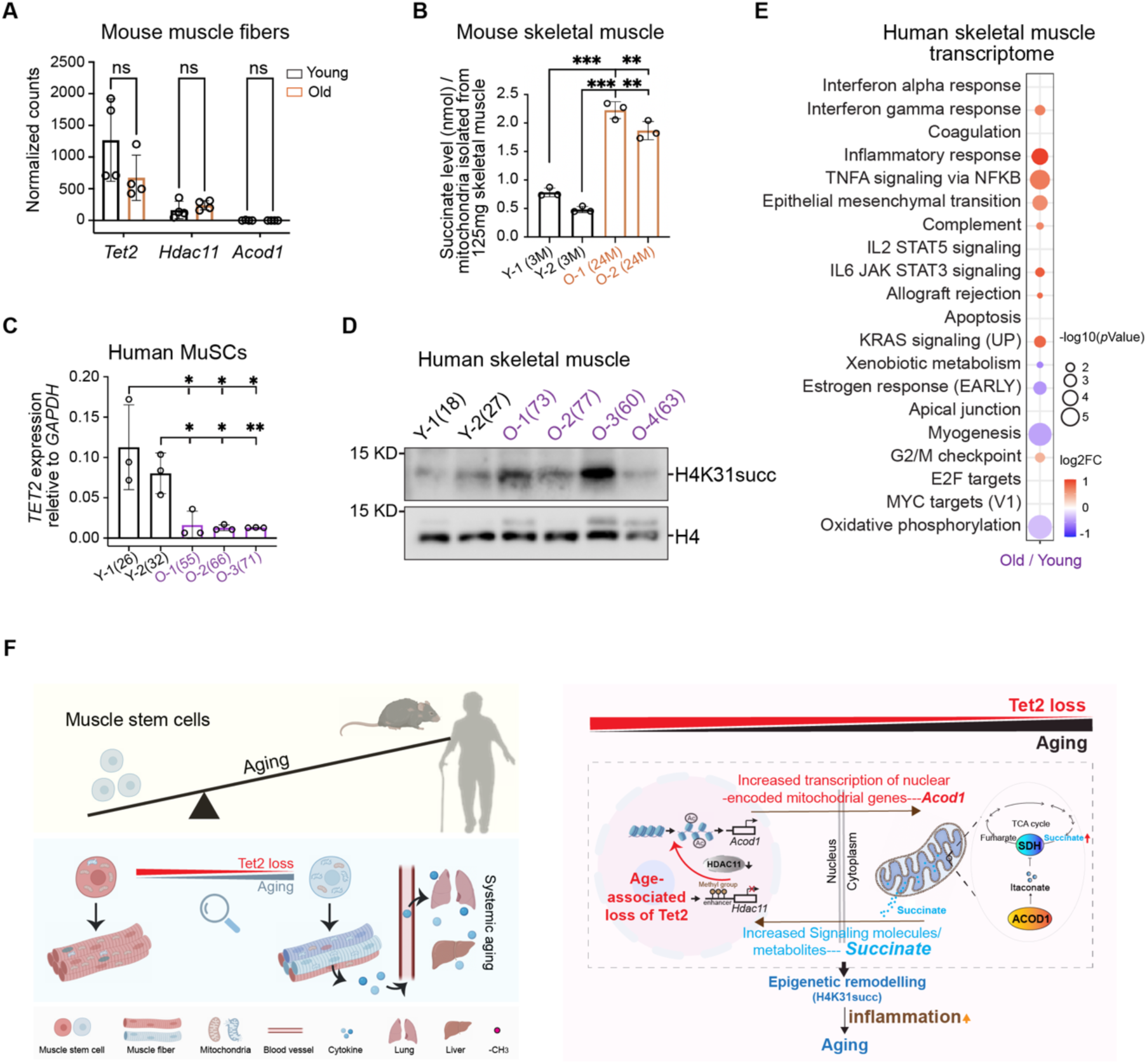
Age-associated decline of *Tet2* expression in MuSCs induces multi-organ inflammaging during aging. Related to Figure 7. **(A)** Normalized counts of *Tet2*, *Hdac11*, and *Acod1* using RNA-seq data obtained from MuSCs-free muscle fibers isolated from young and old mice, respectively (n=4 in each group). RNA-seq data were from GSE193261. **(B)** Absolute succinate content measured by coupled enzyme assays using mitochondria isolated from quadriceps of young (3-month-old) and old (24-month-old) mice, respectively (n=3 in each group). **(C)** Analysis of relative mRNA level of *Tet2* by RT-qPCR using MuSCs isolated from skeletal muscles of young and old people, respectively. The ages of the donors were indicated in the parentheses below the horizontal axis. **(D)** Immunoblotting analysis of H4K31succ in skeletal muscles from young and old people, respectively. H4 served as a loading control. The ages of the donors were indicated in the parentheses. **(E)** Bubble plot comparing functional enrichments of MSigDB (v2023.2) hallmarks for genes differentially expressed in skeletal muscle from young and old people. Dot sizes: - log_10_ (*p* value); dot colors: average log2 fold change (Old/Young) of all genes in each functional term (blue to red). RNA-seq data were from GSE167186. **(F)** Model graph illustrating the multi-organ aging controlled by a small set of MuSCs. **(A)-(C)** Data were shown in mean ± SD form. Data were analyzed by unpaired t-test. ** indicated *p* < 0.01; *** indicated *p* < 0.001; ns indicated not significant.

